# RiboTag-Seq Reveals a Compensatory cAMP Responsive Gene Network in Striatal Microglia Induced by Morphine Withdrawal

**DOI:** 10.1101/2020.02.10.942953

**Authors:** Kevin R. Coffey, Atom J. Lesiak, Russell G. Marx, Emily K. Vo, Gwenn A. Garden, John F. Neumaier

## Abstract

Microglia have recently been implicated in dependence to opioids. To investigate this directly, we used RNA sequencing of ribosome associated RNAs from striatal microglia (RiboTag-Seq) after the induction of morphine tolerance and then the precipitation of withdrawal by naloxone. We detected large, inverse changes in RNA translation following opioid tolerance and withdrawal, and bioinformatics analysis revealed an intriguing upregulation of cAMP-associated genes that are involved in microglial motility, morphology, and interactions with neurons. Three-dimensional histological reconstruction of microglia revealed changes in process branching and termination following opioid tolerance that were rapidly reversed during withdrawal and were consistent with cAMP effects on microglia morphology. Direct activation of Gi-coupled DREADD receptors in microglia, rather than mimicking the effects of morphine, exacerbated opioid withdrawal. Together these indicate that microglial response to cAMP signaling can mitigate the rapid manifestations of opioid withdrawal.

---

**O**pioid abuse has reached epidemic proportions in the United States and is responsible for more than 40,000 overdose deaths each year ^1,2^. The physiological consequences of chronic opioid use include tolerance (which increases the risk for accidental overdose) and withdrawal, which produces physical, emotional, and behavioral symptoms and is a major deterrent to discontinuing of opioids. The symptoms of withdrawal can include anxiety, dysphoria, autonomic instability, muscle and abdominal cramping, diarrhea, and many additional features of severe malaise. While the motivations for continued drug use are complex, the threat of withdrawal provokes anticipatory anxiety and avoidance of treatment seeking, both for the addiction itself as well as for other medical problems. Besides substitution therapy (such as methadone or buprenorphine), currently available treatments for opioid withdrawal are not sufficiently effective and developing new strategies for diminishing withdrawal may offer important health benefits and increase the chances of those suffering from substance abuse choosing to discontinue opioids. Molecular and behavioral responses to opioids have been thought to be mediated primarily by neurons, but there is accumulating evidence that other cell types play a prominent role in drug addiction ^3–5^. For example, microglia are now understood to be partners in plasticity and may provide new therapeutic avenues ^6^.

Little is known about the interface of opioids, microglia, and neuroinflammation, but it is likely that there are both direct and indirect effects of opioid receptor activation on microglia function. Resting microglia reside in the brain parenchyma, engage in synaptic pruning ^7–9^ and in chemical and mechanical surveillance for the presence of challenges such as infection or neuronal damage. Once engaged, microglia may release a wide range of effector molecules and enzymes, phagocytose debris, remodel nearby synapses, etc. ^10^. Opioid administration modulates microglial activity. For example, the density of Iba-1-positive microglia in cortex and striatum are reduced after sub-chronic opioid treatment but increased again after spontaneous withdrawal ^11^. Neuroinflammation also has a complex role in pain and tolerance to opioid analgesics and microglia have been recently identified as playing a key role in the paradoxical development of opioid-induced hyperalgesia, ^12,13^ although there are likely multiple other mechanisms involved. Morphine potentiates lipopolysaccharide-induced microglia activation via both opioid receptor mediated and TLR4-mediated pathways ^14^. Microglia do express mu opioid receptors, ^15,16^ yet mu opioid receptor deletion from microglia does not change opioid tolerance and antinociception ^17^. However, gross microglia inhibition by the compounds minocycline or ibudilast ^18^ or blocking microglial pannexin channels ^19^ produces a dose-dependent reduction in opioid withdrawal, suggesting a mostly indirect effect of microglia on opioid-mediated behaviors.

Both tolerance and withdrawal from morphine are associated with increased microglial activation, but diverse signaling mechanisms may be involved. Opioid withdrawal can intensify pain sensitivity, termed hyperalgesia, and this appears to involve pro-inflammatory processes including activation of TLR2, TLR4, the NLRP3 inflammasome, and NF-KB ^12,14,20–22^, suggesting that several microglial signaling pathways may be involved. Since abused opioids are used cyclically, there is a potential for the repeated exposure to prime or condition microglia such that they produce an exaggerated immune response to other stimuli later, including withdrawal, which has been shown to activate microglial release of cytokines ^11^. While most previous studies of microglia have examined later stages in this process, we hypothesized that microglial function and gene expression are altered at much earlier time points, even during the rapidly induced behavioral changes associated with precipitated withdrawal.

One method for assessing rapidly evolving adaptations to environmental challenges is to investigate alterations in RNA expression and translation. A great deal of microglial RNA translation occurs on ribosomes in fine processes that are not recovered by mechanical methods of collecting and isolating microglia. Indeed, a recent report indicated that FACS sorting of microglia led to significant alterations in immediate early gene expression when compared to RNA isolation using a rapid ribosome pulldown ^23^. Thus, we decided to use the RiboTag strategy to immunopurify microglial ribosome-associated mRNAs, which provides an excellent yield of highly enriched microglial mRNA that were actively undergoing translation, followed by next-gen sequencing of RiboTag mRNA (RiboTag-Seq). Using a well-characterized morphine administration schedule that produces rapid tolerance ^24^ and collected ribosome-associated mRNA from striatal microglia in order to capture the rapid changes in translation in response to naloxone-precipitated withdrawal in mice expressing RiboTag in brain microglia. We provide, to our knowledge, the first complete picture of the translational consequences of severe opioid withdrawal on microglia. We find that RNA actively undergoing translation is markedly altered by opioid tolerance, and many of these changes are rapidly reversed by naloxone-precipitated withdrawal. Intriguingly, many of the RNAs that are altered related to cyclic-AMP signaling, and inhibiting cAMP-mediated signaling in microglia with Gi/o coupled DREADD activation exacerbated the acute behavioral signs of withdrawal. We also observed changes in microglia process branching and termination following opioid tolerance that was rapidly reversed during withdrawal and was consistent with cAMP effects on microglia morphology ^25^. Together, this data suggests that microglia are actively involved in both opioid tolerance and withdrawal in a compensatory manner.

## Results

### Escalating Morphine Causes Locomotor Sensitization while Naloxone Precipitates Withdrawal

Male and female mice were administered “drug”, morphine or saline for six days, then given a “treatment” of naloxone or saline in a 2×2 design yielding 4 groups: saline+saline (SS), saline+naloxone (SN), morphine+saline (MS), and morphine+naloxone (MN) (Figure 1b). Data were analyzed by 2-way ANOVA and ‘Tukey-Kramer’ corrected multiple comparisons. There was a significant main effect of “drug” on locomotion (f(15,1)=10.83, p=.005) and a significant interaction of “drug” and “treatment” (f(15,1)=23.35, p<.01). Morphine-saline treated animals displayed robust locomotor sensitization, traveling significantly more than MN, SN, and SS animals (all**\*** p<.01; Figure 2b). There was a significant main effect of “drug” on body contraction (f(15,1)=6.63, p=.021) and a significant interaction of “drug” and “treatment” (f(15,1)=6.16, p=0.025), with the MN group displaying classic signs of physical discomfort, spending more time hunched and contracted than any other group (all**\*** p<.01; Figure 2c). There was also a significant interaction of “drug” and “treatment” on immobility (f(15,1)=26.28, p=0.0001) where MN animals stayed immobile longer than any other group. (all* p<.01; Figure 2d). SS animals and SN animals did not significantly differ in any tested behavior.

**Figure 1.**
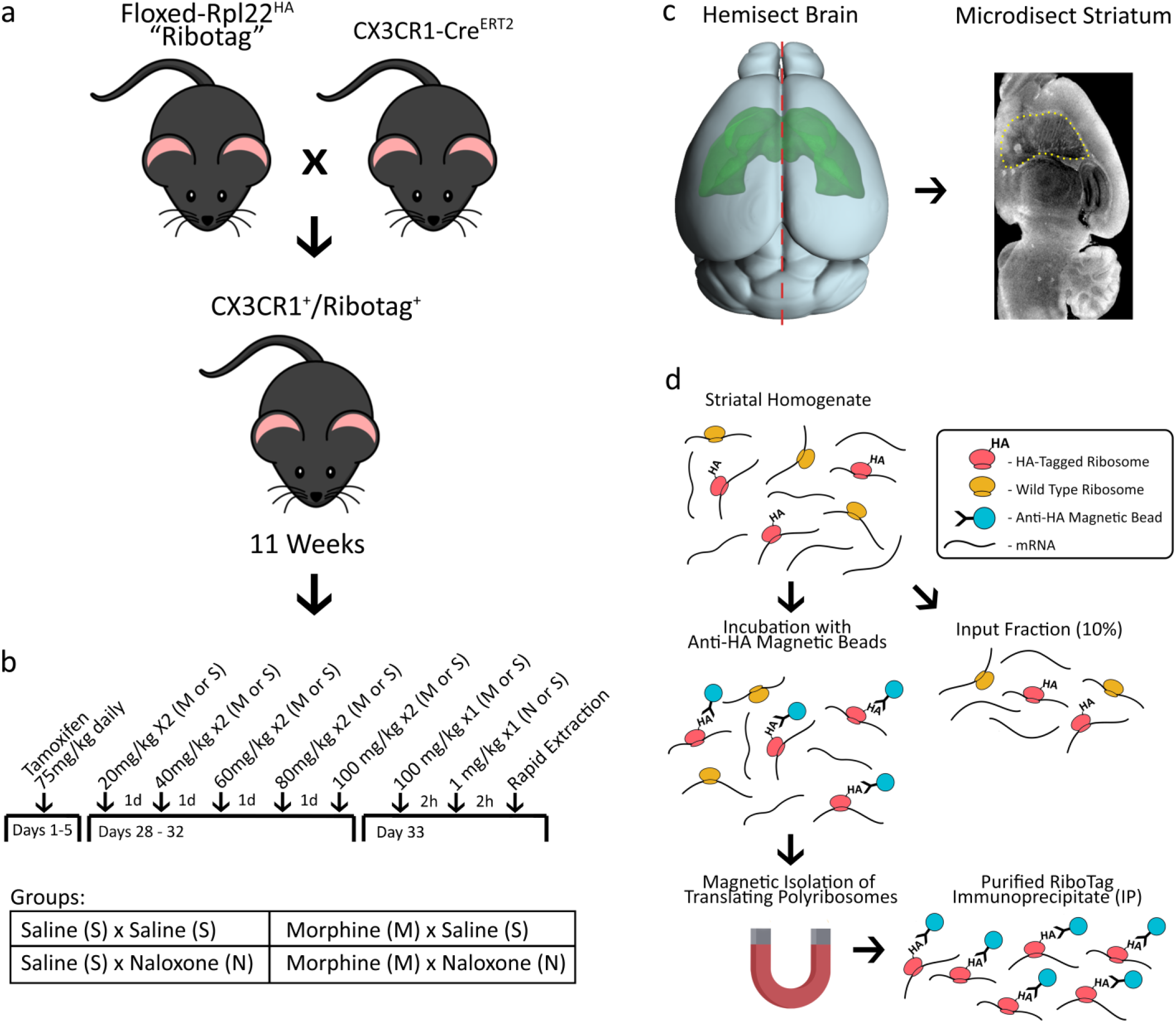
Experimental Overview. **a**, Experimental male and female mice were generated by crossing tamoxifen-inducible CX3CR1-CreERT2 hemizygous mice with homozygous floxed RiboTag mice. **b**, The experiment was performed as a 2×2 treatment design (n=6/group) with all mice receiving tamoxifen for seven days at six weeks. At ~11 weeks of age the mice received either morphine or saline for five days, on a twice daily schedule with the morphine dose escalating. On day 6 the mice received 100 mg/kg *ip* morphine or equivalent volume saline. Then 2 hours later they received either naloxone (1 mg/kg *sc*) or saline. **c**, Brains were rapidly extracted, hemisected, and the striatum was micro-dissected and homogenized for RiboTag. **d**, RiboTag isolation procedure involves incubating tissue RNA with anti-ha magnetic beads and performing magnetic isolation of ribosome bound RNAs.

**Figure 2.**
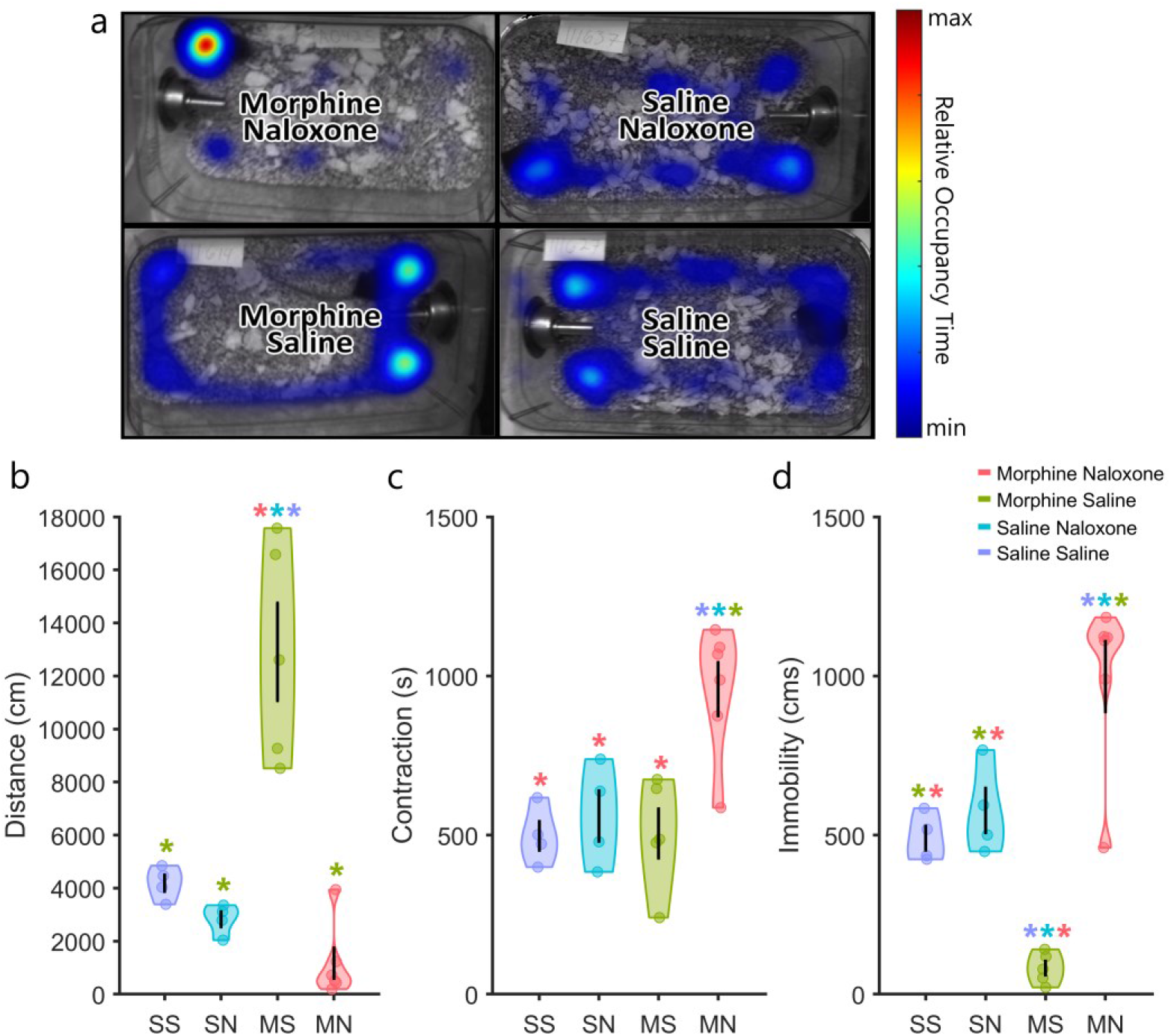
Behavior. **a**, Automatic scoring of morphine and withdrawal behavior. **b**, Morphine administration induces locomotor sensitization while naloxone precipitated morphine withdrawal induces **c**, body hunching (contraction) and **d**, immobility. Saline-saline animals and saline-naloxone animals did not significantly differ in any tested behavior.

### Immunoprecipitation and Sequencing Descriptive Statistics

Tissue homogenate was spun down and the supernatant was split, 90% for immunoprecipitation and 10% for Input (Figure 1d). RNA extraction from Input samples yielded 120 ±11 ng (mean±sem) of RNA while IP samples yielded 51±4 ng (mean±sem) of RNA. 10ng of RNA from each sample were used for library preparation. RNA sequencing from Input samples yielded 4.09±0.476 million reads (mean±sem) while IP samples yielded 2.78±0.241 million reads (mean±sem). All fastq files passed FastQC basic statistics, per base quality, and per sequence quality scores.

### Microglia Specific Markers Are Highly Enriched in RiboTag-Seq

Differential expression analysis of IP vs Input samples showed dramatic enrichment of microglia specific RNAs (Figure 3a). 10,540 genes were significantly enriched in the IP samples. Among the top 25 most enriched genes are many classic microglia markers such as C1qa, Tmem119, Itgam (CD11B), and CX3CR1 (Figure 3a). Gene set enrichment analysis of IP vs Input samples demonstrated enrichment of classic microglia pathways, such as cytokine binding, purinergic receptor activity, and NF-kB binding (Figure 3b). All of the data and analyses from differential expression and GSEA for enrichment is available in the Supplementary Code and Data..

**Figure 3.**
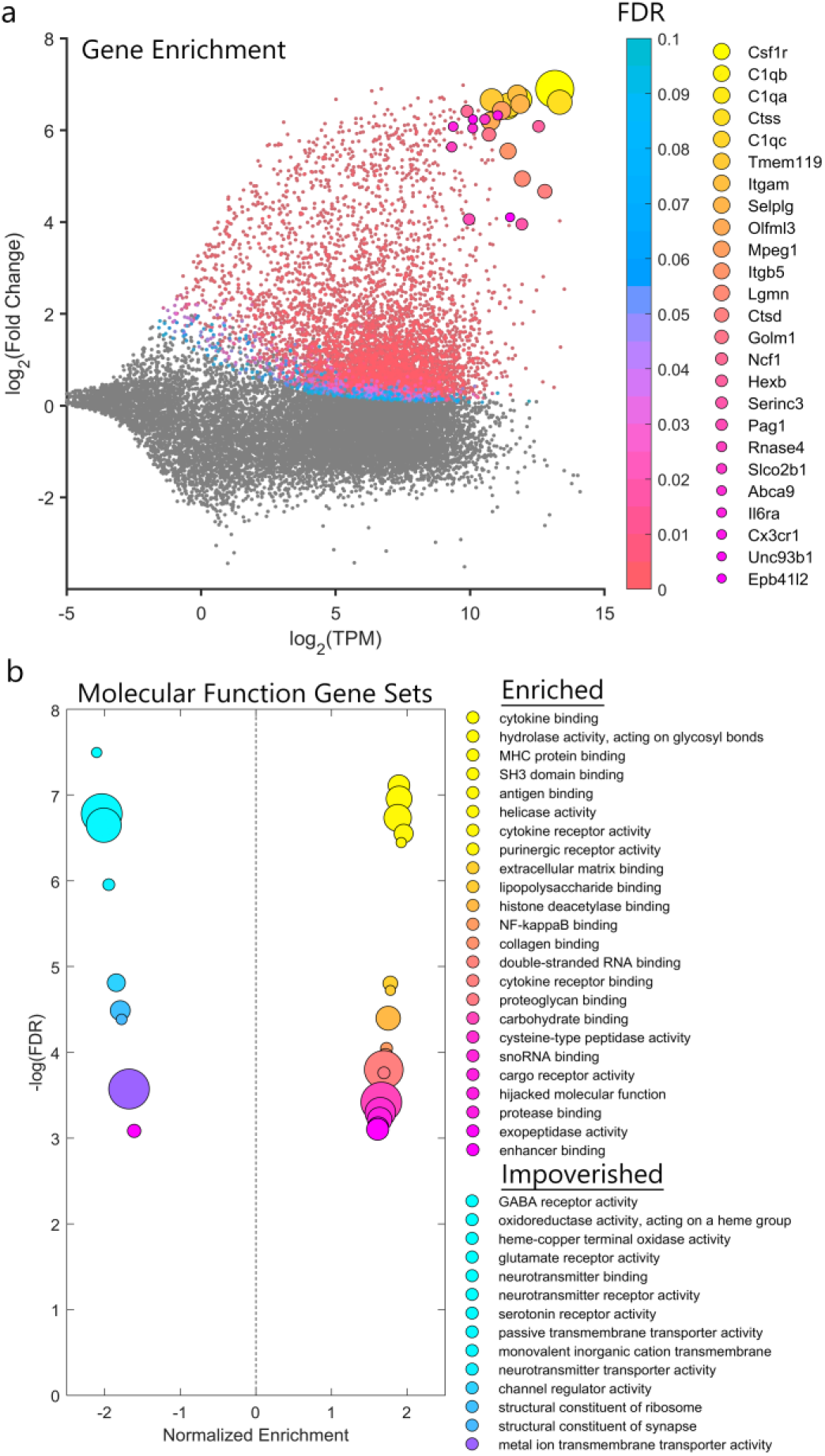
RiboTag Enrichment. **a**, The RiboTag-Seq samples are dramatically enriched with microglial markers such as Tmem119 and CX3CR1. **b**, GSEA reveals an upregulation of genes associated with classical microglial gene sets such as “cytokine binding” and “purinergic receptor activity”.

### Morphine Tolerance and Withdrawal Produce Inverse Patterns of Differential Gene Expression

Naloxone administration in non-morphine-dependent animals yielded very few differentially expressed genes (DEGs) in microglia or the striatum as a whole (SN vs SS; Figure 4a,d). However, escalating morphine exposure produced large changes in both microglia (MS IP vs SS IP; Figure 4b) and in the striatum as whole (MS Input vs SS Input; Figure 4e). In both the IP and Input samples, morphine tolerance produced similar numbers of upregulated and downregulated genes. In the microglia (MS IP vs SS IP), morphine tolerance produced 555 upregulated and 653 downregulated genes. In the striatum as a whole (MS Input vs SS Input), morphine tolerance produced 1207 upregulated and 1372 downregulated genes. By contrast, withdrawal produced a dramatic upregulation in genes with roughly half as many downregulated. In the microglia (MN IP vs MS IP), morphine tolerance upregulated 706 and downregulated 344 genes. In the striatum as a whole (MN Input vs MS Input), morphine tolerance upregulated 1373 and downregulated 910 genes. In both the IP and Input samples there was a significant inverse relationship between gene expression changes during morphine tolerance and withdrawal. (IP: R^2^=0.86, p<.001; Figure 4g; Input: R^2^=0.90, p<.001; Figure 4h). Genes that were downregulated during morphine tolerance tend to be upregulated during withdrawal, and vice-versa.

**Figure 4.**
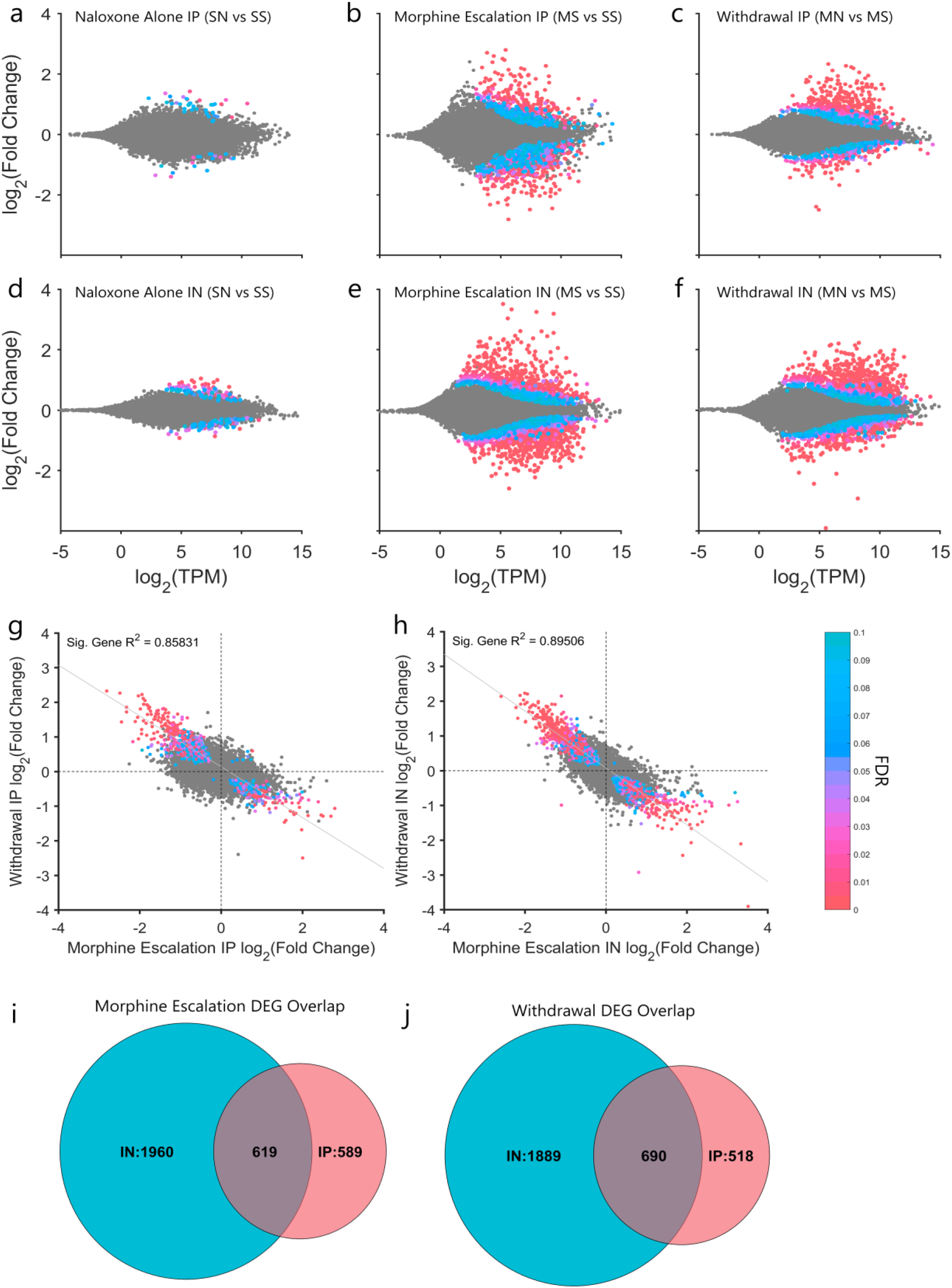
Differential Expression Analysis. Naloxone administration to non-morphine-dependent animals has little effect on gene expression in the **a**, IP samples or **d**, Input samples. Morphine tolerance produces roughly equal numbers of upregulated and downregulated DEGs in both the **b**, IP sample and **e**, Input sample. Naloxone administration to morphine dependent animals produces mainly upregulation of DEGs in both the **c**, IP and **f**, Input samples. Gene expression changes during withdrawal are inversely correlated to gene expression changes during morphine tolerance in both the **g**, IP sample and **h**, Input sample. Roughly half of IP DEGs were also differentially expressed in the Input samples for both **i**, morphine tolerance and **j**, withdrawal, suggesting a common mechanism of action in neurons and microglia.

There was substantial overlap in IP and Input DEGs for morphine tolerance (619; Figure 4i) and withdrawal (690; Figure 4j). This may be due to a common mechanisms in microglia and other cell types as well as the fact that microglia make up around 10% of the cells in the striatum ^26^. Still, there are many DEGs specific to the Input and IP samples. All of these genes and their corresponding statistics can be accessed in the Supplementary Code and Data.

### Gene Set Enrichment Analysis of IP and Input RNA during Morphine Tolerance and Withdrawal

GSEA is a method of deriving biological meaning from differential expression data based on previously annotated data sets. Genes sets generated from experimental data are compared to differential expression data to determine if the distribution of differential expression within a gene set is shifted toward upregulation or downregulation. For the IP samples, we found that morphine tolerance was associated with upregulation of gene sets related to protein folding and the unfolded protein response (UPR; Figure 5a), responses associated with cellular stress. Similar genes associated with the heat shock protein (HSP90) chaperone cycle were also upregulated (Figure 5b). Morphine tolerance was also related to downregulation of genes commonly associated with dendrite development and synaptic organization (Figure 5a).

**Figure 5.**
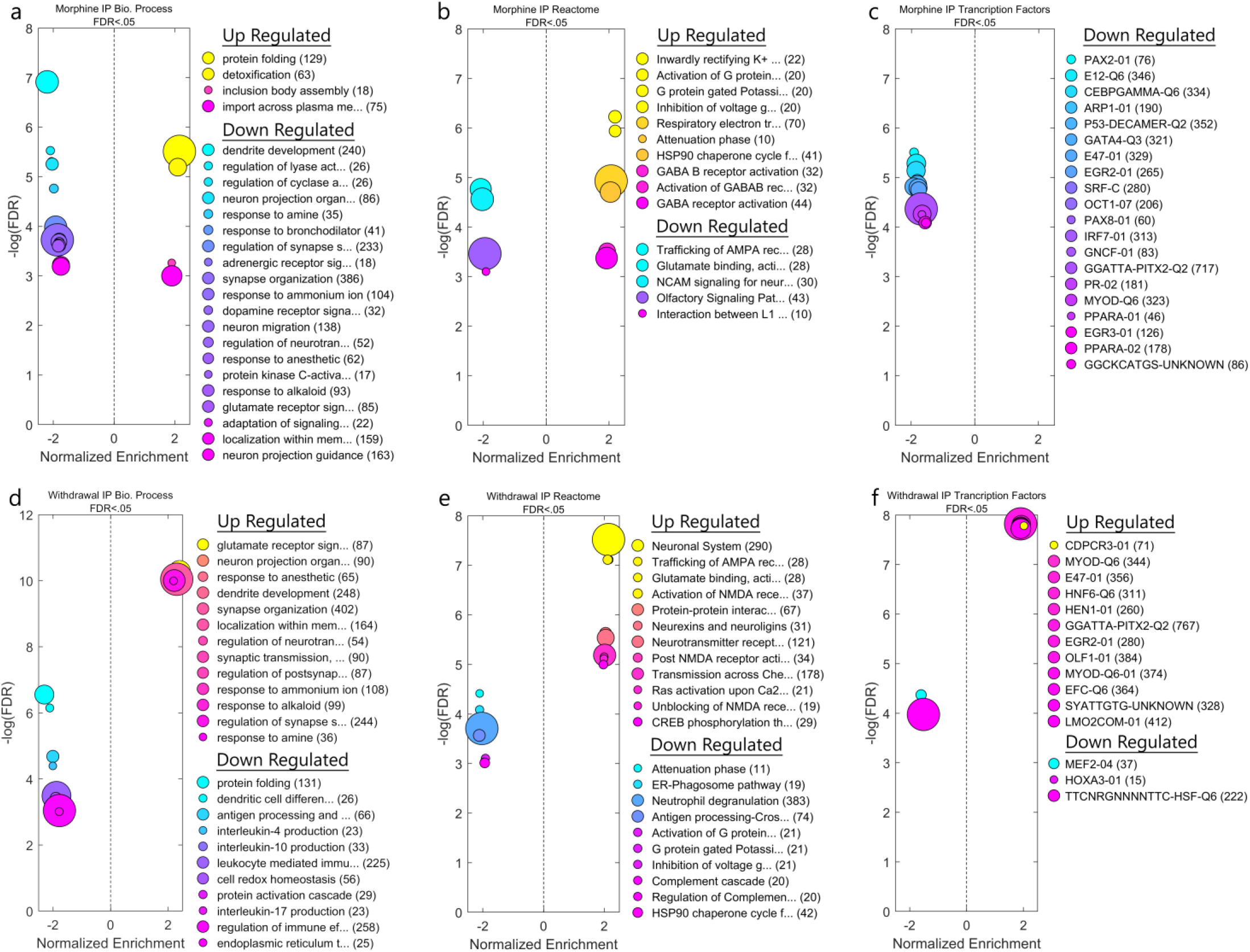
Gene Set Enrichment Analysis. GSEA reveals many annotated gene sets that are significantly upregulated or downregulated in microglia. **a,b,c**, For example, morphine tolerance upregulates protein folding genes and inwardly rectifying K+ channels, and downregulates synaptic organization genes and genes associated with the transcription factor PAX2. **d,e,f**, On the other hand, withdrawal upregulates synaptic organization genes, CREB phosphorylation, and genes associated to the immune related transcription factor EGR2, and downregulation of interleukin production, and G protein gated K+ channels.

Naloxone-precipitated withdrawal produced roughly opposite changes in IP gene set enrichment. Gene sets associated with synaptic organization were upregulated (Figure 5d, e), while protein folding gene sets were downregulated (Figure 5d, e). Several transcription factor targets were also found to be enriched (figure 5f), which may be viable targets for manipulation of entire gene networks. This analysis was completed for the Input samples as well and are shown in Supplementary Figure 1. The entire GSEA pipeline for IP, Input, and Enrichment groups can be found in the Supplementary Code and Data.

### WGCNA Reveals a cAMP Responsive Gene Network in Striatal Microglia induced by Morphine Withdrawal

Weighted gene co-expression network analysis (WGCNA) ^27^ was used to produce a signed topological overlap matrix representing the network connectedness of RiboTag-Seq expression across all samples. This matrix was transformed into a hierarchical tree and clustered with the dynamic tree cut library for R ^28^. Clusters produced from this procedure are referred to as gene modules, and randomly re-named using the “Crayola Color Palette” in order to combat attribution of meaning or importance to numbered modules. Modules were merged by clustering on module eigengenes with a cut height of 0.85. All details of clustering can be found in the WGNA R and MATLAB codes in the Supplementary Code and Data. A minimum spanning tree of the entire RiboTag-Seq matrix and corresponding module labels were projected into 2D space using a force directed layout (Figure 5a). Significance values for morphine tolerance (Figure 5b) and withdrawal (Figure 5c) are overlaid on the minimum spanning tree to visualize the relationship between modules and gene significance. Highly significant DEGs form networks that are sometimes overlapping for morphine tolerance and withdrawal, whereas others were unique to one condition or the other.

Of particular interest are modules that were differentially expressed between treatment groups. The “Sheen Green” module and “Timberwolf” modules contain genes which were, on average, downregulated during morphine tolerance (Supplementary Figure 2a), and upregulated during withdrawal (Supplementary Figure 2b). These modules were analyzed thoroughly to determine if network membership was meaningful or just capturing a simple inverse relationship in differential expression. Undirected network graphs were generated for both the Sheen Green module (Figure 6a) and the Timberwolf module (Figure 6d) and were projected into 2D space using a circle layout. The benefit of a circle layout over a traditional force layout is an increase in legibility. This method, along with many others, is shared in our @WGCNA MATLAB class available at https://GitHub.com/DrCoffey/WGCNA. Each gene’s importance within the network is defined using a centrality metric. Centrality was calculated using “closeness” for the distance calculation, and the sum of each node’s edge weight (similarity) as the “cost” function. Genes that are central to the network form many strong connections with other genes, which is represented by many bright lines connecting a gene to others in the circle graphs (Figure 7a, d). The top four most central genes in each module are highlighted with an orange dot.

**Figure 6.**
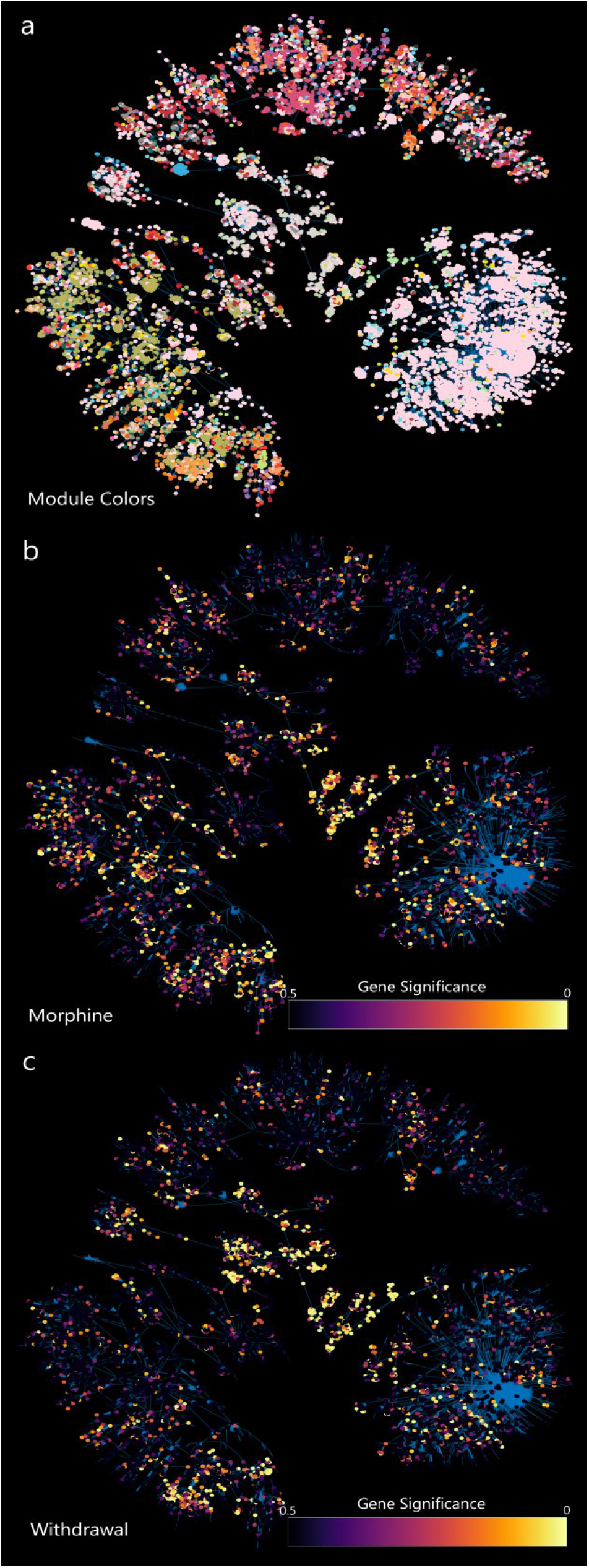
WGCNA Minimum Spanning Tree. **a**, Genes are coded by module color and projected into a 2D minimum spanning tree using a “force” layout. **b**, The same genes are coded by differential expression (q-value) data during morphine tolerance. **c**, The same genes are coded by differential expression (q-value) data during morphine withdrawal. Functionally related genes tend cluster in modules, and some are related to both morphine and withdrawal while others are uniquely effected by one, the other, or neither.

**Figure 7.**
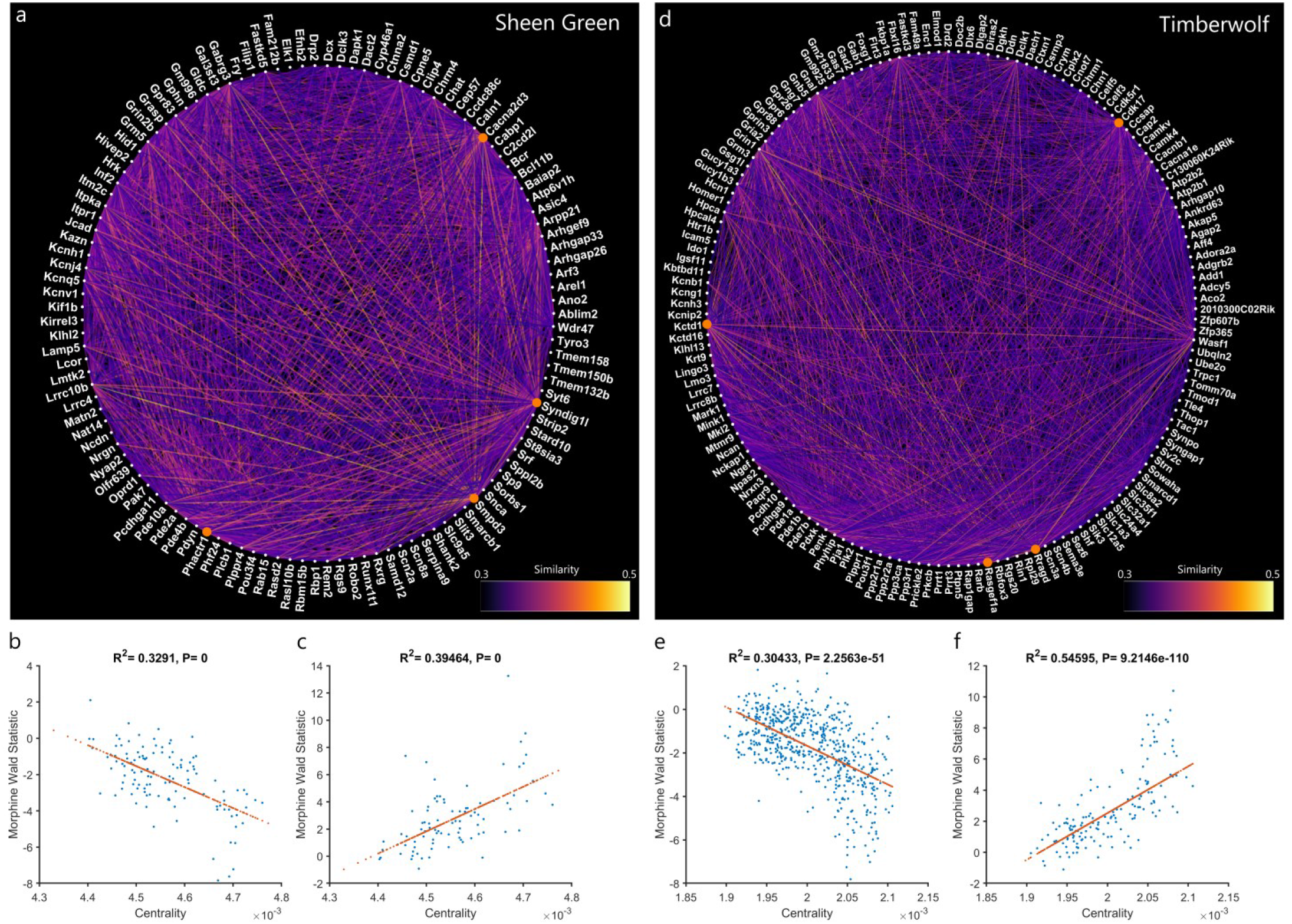
cAMP Related Gene Modules. **a,d**, Gene modules projected into 2D space using a circle layout. Line brightness represents gene similarity as calculated by WGCNA. The top 4 most central genes in each network are labeled with an orange dot. Centrality (a measure of network importance) in the “Sheen Green” network is highly correlated to differential expression for both **b**, morphine tolerance and **c**, withdrawal. Centrality in the “Timberwolf” network is also highly correlated to differential expression for both **e**, morphine tolerance and **f**, withdrawal. In both cases morphine tolerance suppresses central genes and withdrawal induces them.

For both modules, network centrality was highly correlated to gene significance for morphine tolerance and withdrawal. For the Sheen Green module, network centrality was negatively correlated to morphine tolerance Wald-statistics (IP MS vs IP SS; Figure 7b), while network centrality was positively correlated to withdrawal Wald-statistics (IP MS vs IP SS; Figure 7c). For the Timberwolf module, network centrality was negatively correlated to morphine tolerance Wald-statistics (IP MS vs IP SS; Figure 7e), while network centrality was positively correlated to withdrawal Wald-statistics (IP MS vs IP SS; Figure 7e). These strong relationships between centrality and differential expression suggest that network membership is related to the effects of morphine tolerance and naloxone-precipitated withdrawal.

Many of the genes within these networks are downstream targets of cAMP signaling and promote reorganization of synapses and the actin cytoskeleton. The fact that unbiased clustering with WGCNA identified functionally related gene networks suggests that it is an extremely useful tool for discovery of novel genes of interest, independent of any historical biases associated with gene ontology analyses. Other gene modules may also be of interest but are not analyzed deeply in this report and can be reviewed in the Supplementary Code and Data. The same analytic scheme was also performed on the Input RNA representing the total striatal transcriptome.

### Activation of hM_4_Di in Microglia worsens Symptoms of Withdrawal

Given the discovery of these cAMP signaling associated gene modules, we decided to test whether manipulating cAMP signaling in microglia altered withdrawal-associated behaviors. CX3CR1-hM_4_Di animals and CX3CR1-CreERT2 controls were subject to our escalating morphine procedure, followed by naloxone precipitated withdrawal as described in the methods. Twenty minutes before naloxone administration, CX3CR1-hM_4_Di animals received CNO (3mg/kg *ip*) or vehicle, and CX3CR1-Cre^ERT2^ controls received CNO (3mg/kg *ip*) 20 min before naloxone. CX3CR1-hM_4_Di animals who received CNO showed a non-significant trend toward reduced distance traveled as compared to CX3CR1-hM_4_Di animals who received Vehicle, and CX3CR1-CreERT2 controls who received CNO (Figure 8a). CX3CR1-hM_4_Di animals that received CNO displayed significantly more body contraction as compared to vehicle treated CX3CR1-hM_4_Di animals (t(10)=2.35, p = 0.046; Figure 8b), and CX3CR1-Cre^ERT2^ controls who received CNO (t(10)=2.50, p = 0.030; Figure 8b). Further, CX3CR1-hM_4_Di animals who received CNO also displayed significantly more immobility as compared to CX3CR1-hM_4_Di animals who received Vehicle (t(10)=2.68, p = 0.023; Figure 8c), and CX3CR1-Cre^ERT2^ controls who received CNO(t(10)=2.17, p = 0.050; Figure 8c). CX3CR1-hM_4_Di animals who received vehicle and CX3CR1-Cre^ERT2^ controls who received CNO did not differ significantly on any measure. Taken together, these results suggest that activation of hM_4_Di and reduction in cAMP specifically in microglia exacerbated the signs of withdrawal.

**Figure 8.**
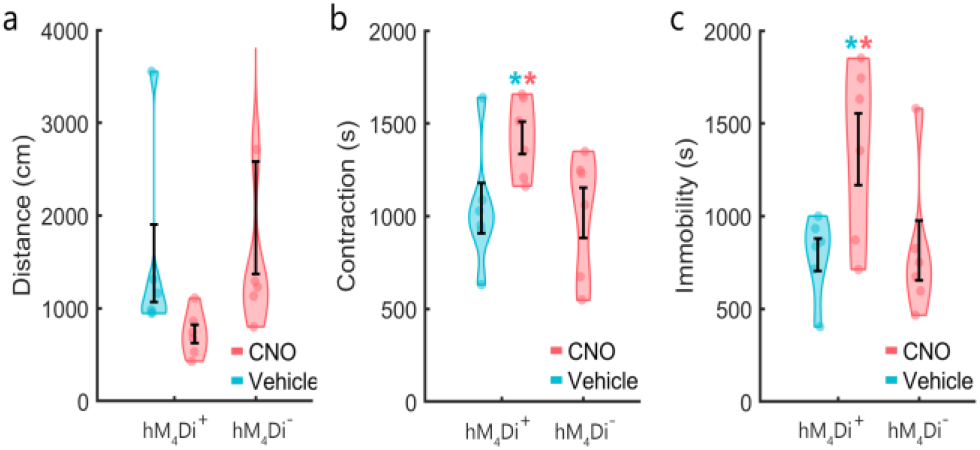
Activation of hM_4_Di in Microglia Exacerbates Withdrawal Symptoms. **a**, CX3CR1-hM_4_Di animals who received CNO showed a non-significant trend toward reduced locomotion. **b**, CX3CR1-hM_4_Di animals that received CNO displayed significantly more body contraction as compared to vehicle treated CX3CR1-hM_4_Di animals and CX3CR1-CreERT2 controls who received CNO. **c**, CX3CR1-hM_4_Di animals who received CNO also displayed significantly more immobility as compared to CX3CR1-hM_4_Di animals who received vehicle and CX3CR1-Cre^ERT2^ controls who received CNO. CX3CR1-hM_4_Di animals who received vehicle and CX3CR1-CreERT2 controls who received CNO did not differ significantly on any measure.

### Striatal Microglia Morphology is Altered Morphine Tolerance and Withdrawal

A separate group of 16 wild type animals (8 male, 8 female) underwent identical morphine administration and naloxone precipitated withdrawal using the same 2×2 design as the RiboTag-Seq animals. At sacrifice, animals were perfused, sectioned at 40µm, and stained for Iba1. Confocal Images were acquired at 40x (Figure 9a) and individual microglia were extracted (Figure 9b) and analyzed in 3 dimensions using 3DMorph software ^29^. Data were analyzed by 2-way ANOVA and ‘Tukey-Kramer’ corrected multiple comparisons. There was a significant main effect of “drug” on volume (f(243,1)=6.68, p=.010). Morphine-saline microglia showed a slight increase in volume over saline-naloxone animals (p<.01; Figure 9c,f). There was a significant main effect of “drug” on branch points (f(243,1)=16.91, p<0.001) and a significant interaction of “drug” and “treatment” (f(243,1)=10.4, p=0.0014), with the morphine-saline microglia displaying an increase in branching, while naloxone rapidly reversed this effect (all p<.01; Figure 9d,g). There was a significant main effect of “drug” on process termination (end points: f(243,1)=18.39, p<0.001) and a significant interaction of “drug” and “treatment” (f(243,1)=11.6, p=0.0008), with the morphine-saline microglia displaying and increase in end points, while naloxone rapidly reversed this effect (all p<.01; Figure 9e,h). SS and SN microglia did not significantly differ in any morphological metric.

**Figure 9.**
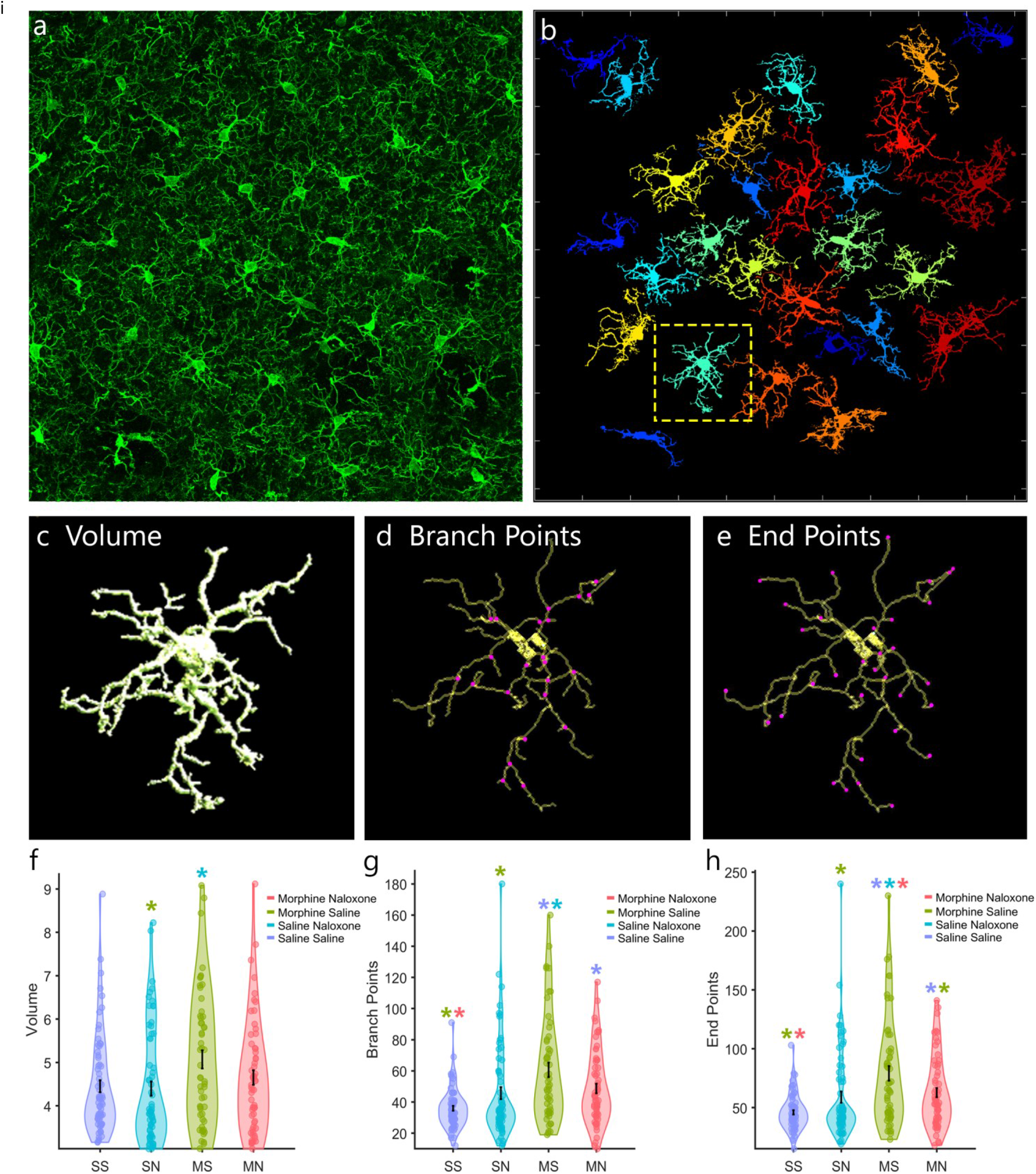
Morphine Tolerance and Withdrawal Inversely Effects Microglia Morphology. **a**, Maximum intensity projection of striatal microglia captured at 40x (30µm thick). **b**, Automated cell segmentation and removal of incomplete (border touching) cells using 3DMorph. **c**, 3D volumetric reconstruction of the highlighted microglia in the above panel. **d**, Automatically calculated branching points generated from a 3D skeletonization of the cell. **e**, Automatically calculated end points generated from a 3D skeletonization of the cell. **f**, There was a main effect of morphine administration on increasing cell volume. **g**, There was a significant main effect of morphine on process branching, and a significant drug*treatment interaction. After correction for multiple comparisons there was only a trend for naloxone to reverse the increase in branching. **h**, There was a significant main effect of morphine on process terminations (end points), and a significant drug*treatment interaction. Morphine tolerance significantly increases the number of microglial process end points, while withdrawal reversed this effect and reduced end points back to baseline. Saline-saline animals and saline-naloxone microglia did not significantly differ in any morphological metric.

## Discussion

Microglia are the innate immune cells of the brain but are also involved in intercellular interactions that are not directly related to immune defenses. The processes and filopodia of “resting” microglia constantly engage in surveillance of the parenchyma and are critical for cell-cell interaction, sensing of chemical cues, and are involved in synapse formation and elimination ^8,9,30,31^. Microglia have been described as quiescent at baseline, but their interactions with nearby neurons and astrocytes are now recognized to play important roles in the maintenance of normal brain function ^7,9,32^. Upon priming and activation, microglia undergo dramatic morphological changes when they may withdraw their highly ramified processes, migrate to the sites of infections or injury in order to destroy pathogens, and remove damaged cells. However, in this report we identify dynamic changes in microglia that occur in advance of activation into states that are generally associated with neuroinflammation.

### The Dynamic Nature of Microglia is Ideally Suited to RiboTag Analysis

Recent work has shown that microglia play a role in the sequelae of opioid withdrawal ^4,5,18,19^, but the extent and nature of their compensatory and/or deleterious actions is still unknown. Microglia are highly dynamic cells that respond to changes require rapid protein production that may relate to changes in transcription but often involve differential loading of RNAs onto ribosomes that is regulated by non-transcriptional mechanisms. RiboTag-Seq allows the capture of a complete snapshot of the translatome with precise timing. In this report we take full advantage of the consistent, time-locked nature of naloxone-precipitated withdrawal to map the translatome of striatal microglia during their initial response to the unique insult of opioid withdrawal. Through a combination of guided (GSEA) and unguided (WGCNA) bioinformatics strategies, we have discovered several gene sets, transcription factors, and gene networks that may prove to be valuable targets for novel treatment of opioid withdrawal.

### Computational Removal of Input Contamination Improves the Reliability and Interpretability of RiboTag-Seq Analysis

Initially, RiboTag-Seq studies relied on the concept of “enrichment” to determine which genes are specific to the cell type of interest and are positively differentially expressed in the IP samples versus the Input samples. This methodology is extremely effective at isolating genes that are uniquely expressed in a given cell type, as exemplified by Figure 3. Most of the highly enriched genes are microglia exclusive markers, including CX3CR1. However, this technique should not be used to rule out a gene as existing and functional in a cell of interest. First, just because a gene is expressed at lower levels in a particular cell type than in the surrounding tissue does not mean that the gene was detected spuriously, and it certainly does not mean that the action of that gene is not important. Further, direct comparison of IP and Input RNA suffers from a problem known as composition bias. Because RNA-Seq utilizes probabilistic sampling from a finite pool of RNA, the values for gene expression do not represent any absolute quantity. Rather, they represent a proportion of the finite gene pool that is attributed to a gene. The unavoidable consequence of this sampling technique is that the diversity of the finite gene pool directly affects gene quantification measures such as TPM. That is, less diverse gene pools, such as from a unique cell type will inherently raise the proportion of reads mapped to any particular gene.

To avoid this issue, we have developed a new approach to remove Input contamination from IP samples computationally and to determine which gene changes can reliably be ascribed to microglia. To show the validity and reliability of this approach, we modeled a sequencing run of IP and Input samples generated from a known distribution of genes (our finite gene pool). Before modeling we introduce contaminating Input RNA (*gamma generated*) into the IP gene pool (*gamma*). After modeling sequencing we tested our contamination removal method and compared the results to a modeled run without contamination. This technique improves the correlation between estimated TPM and “true” reads, even when the estimate of contamination is approximated (over- or under-estimated). This modeled sequencing run is described in the Methods, and is presented in an interactive MATLAB Live Script in the Supplementary Code and Data, as well as a static HTML file (Supplementary Method 1).

### Morphine Tolerance and Withdrawal Produce Inverse Impacts on the Microglial Translatome

Simple differential expression analysis revealed a striking inverse correlation between morphine- and withdrawal-induced changes in the microglia translatome. Microglia express mu- and delta-opioid receptors and can respond directly to opioid receptor activation caused by morphine and its competitive antagonism by naloxone. However, the broad range of these changes suggests indirect impacts of opioid withdrawal as well. Microglia interact intimately with neuronal synapses and are essential for healthy synaptic formation, pruning, and plasticity ^8,30,31^. Morphine administration and withdrawal are known to inversely affect long term depression ^33^ and have myriad other effects on synaptic activity ^34–36^. It is no surprise then that microglia, which interact with synapses, also show dynamic changes in gene expression.

### Morphine Tolerance Induces the Unfolded Protein Response in Striatal Microglia, While Morphine Withdrawal Induces Synaptic Organization Genes

Morphine tolerance appears to induce a cellular stress effect in microglia as indicated by upregulation of genes that respond to misfolded proteins. Heat shock proteins and chaperone proteins are overrepresented among differentially upregulated genes. The unfolded protein response is a cellular response to stress that is initiated by increased detection of misfolded proteins in the endoplasmic reticulum; the UPR can be a homeostatic response to return the system to baseline but may also lead to stress-induced apoptotic cell death ^37^. Previous studies have detected activation of the unfolded protein response in neurons and astrocytes as well ^38,39^. One interpretation of these results is that despite the appearance of quiescence, repeated morphine exposure primes microglia, preparing them for vigorous response to subsequent challenges.

GSEA also revealed an inverse pattern of regulation among synaptic organization gene sets during morphine tolerance and withdrawal. Morphine tolerance tended to downregulate synaptic organization genes, while withdrawal upregulated them. While many of these gene sets associated with synaptic processes relate to neurons, microglia play an essential role in synapse formation, maintenance, and elimination ^8,30,31^ and they also express genes commonly associated with synapse differentiation such as Syndig1 ^40^. GSEA is explicitly built upon previously interpreted gene associations and many gene sets were originally annotated from transcriptomic studies that used whole tissue homogenates containing glial cells along with neurons, but they may have been interpreted as emanating from neurons. Our conservative computational strategy for eliminating Input contamination from our IP samples increases confidence that these genes are from microglial ribosomes, and are likely to be regulated and functionally important during morphine tolerance and withdrawal.

### WGCNA Reveals a cAMP Responsive Gene Network in Striatal Microglia Inhibited by Morphine Tolerance and Induced by Morphine Withdrawal

WGCNA is an unbiased method of clustering functionally related genes and inferring their network connections. Its use in the present dataset revealed two gene networks that were inversely regulated by morphine tolerance and withdrawal. These networks include numerous downstream targets of cAMP signaling, cytoarchitectural adaptation, and modifiers of synaptic organization. While there are many genes in each module, there was a strong association between the genes’ centrality to the network and their inverse regulation by morphine tolerance and naloxone-precipitated withdrawal. Centrality in an undirected network is analogous to consideration of genes as “hubs” in a directed network. The top 10 most central genes from the Sheen Green and Timberwolf modules are briefly described in Tables 1 and 2. All genes from all modules can be explored in the Supplementary Code and Data. Each of these genes has not previously been associated with opioid tolerance or withdrawal yet has the potential to play important roles in mounting a cellular response that might involve interactions with nearby neurons. These genes include SYNDIG1 (Synapse Differentiation Inducing 1), which has been shown to be expressed in microglia and is involved in synapse organization ^40^, PHACTR1 (Phosphatase And Actin Regulator 1) a multifunctional enzyme with diverse roles, including regulation of synaptic activity and dendritic morphology, and cAMP induced actin remodeling ^41^. Since opioid tolerance and withdrawal induce differential effects of synaptic activity ^33^, it is noteworthy that these two gene modules in general, and the more central genes in particular, are inversely regulated during opioid tolerance and withdrawal.

It is not clear how the microglia might be detecting changes in neuronal excitation, but several receptors on microglia are known to respond to neuronal activity, such as CX3CR1 (the fractalkine receptor) and P2Y12, a receptor that responds to extracellular ATP/ADP. ATP release has also been implicated in the role of microglia and opioid withdrawal, and genetic deletion or inhibition of pannexin-1, an ATP channel, from microglia reduced the signs of opioid withdrawal ^19^. Interestingly, both CX3CR1 and P2Y12 are Gi/o-coupled receptors that inhibit cAMP accumulation in addition to non-canonical signaling effects. Thus, it is possible that altered cAMP signaling in microglia is an important consequence of opioid withdrawal that is not directly mediated by opioid receptors. Indeed, the central role of microglia in opioid-associated hyperalgesia and other types of neuropathic pain has been argued to be independent of microglial mu opioid receptors ^17^ and may involve P2Y12 receptors ^42–44^. While several G protein-gated inwardly rectifying potassium (GIRK) channels were upregulated by morphine, they were downregulated by withdrawal. These are frequently activated by Gi/o-coupled receptors via the release of G_ßγ_ subunits and are implicated in microglial activation ^45^. This downregulation of GIRK translation may be a compensatory response to their over activation during withdrawal. In addition to the numerous downstream targets of cAMP signaling, several phosphodiesterases were among the most strongly upregulated genes by naloxone-precipitated withdrawal. Together, this suggests that cAMP signaling in microglia has an important role in the microglial response to the initial stages of opioid withdrawal.

### Striatal Microglia Morphology is Altered by Morphine Tolerance and Withdrawal

The microglia imaged in this report do not appear to be classically activated, but rather show modest changes in morphology and process branching and termination. While we did not predict these results, they are entirely consistent with a recent report showing that microglia engage in nanoscale surveillance of the brain via cAMP regulated filopodia ^25^. That report showed that decreasing intracellular cAMP results in microglial process elongation, while increasing cAMP causes process retraction. We found that two gene networks discovered in an unbiased analysis suggest that cAMP signaling in microglia is downregulated during morphine tolerance and upregulated during withdrawal. These changes could be involved in the regulation of actin cytoskeleton dynamics that alter microglial process extension and retraction. The morphological changes identified in this report are entirely consistent with these changes in gene translation. Morphine withdrawal is also associated with reduced dendritic spine density and synaptic reorganization in the striatum ^46,47^. Since it is becoming increasingly clear that microglia participate in spine plasticity, we believe that these translational gene networks in microglia may be involved in this phenomenon. RiboTag is sensitive to changes in translation that may relate directly to current challenges that the microglia are responding to, making it very sensitive to detecting genes involved mechanistically in these responses

### cAMP Signaling in Striatal Microglia During Withdrawal May Mitigate Opioid Withdrawal

The induction of cAMP-regulated genes in response to naloxone inhibition of mu opioid receptors seemed counterintuitive to us, since naloxone *decreases* Gi/o signaling in cells that express mu and delta opioid receptors (including microglia). This led us to test this notion directly and investigate the impact of selectively reducing cAMP signaling in microglia by expressing a Gi/o-coupled DREADD receptor ^48^ selectively in microglia, so that we could reduce cAMP chemogenetically, specifically in microglia. The hM_4_Di DREADD receptor activates Gi/o-coupled signaling including inhibition of adenylyl cyclase and has been used in microglia previously ^12^. We found that pretreatment of these mice with CNO exacerbated the hunched body posture and immobility induced by opioid withdrawal. These results suggest that, during the initial instances of withdrawal, cAMP signaling in microglia mitigates the acute behavioral effects of opioid withdrawal, and that the increased translation of cAMP-associated genes may produce compensatory effects that protect against opioid withdrawal. An alternative interpretation is that some combination of inhibiting canonical cAMP-mediated and activating non-canonical signaling by Gi/o-coupled receptors contributes to withdrawal, placing microglia in the center of the opioid withdrawal syndrome. In either case, it seems that upregulation of cAMP-associated genes in microglia does not represent a direct counter-regulatory adaptation to sudden inhibition of opioid receptors located on microglia, but may be an indirect response to adaptations in other cell types such as neurons. Future studies will need to sort out the impact of early vs. later microglial effects on withdrawal, which has many impacts including the acute physiological withdrawal syndrome to changes in motivation that perpetuate opioid abuse.

In summary, we used a new bioinformatics pipeline that shows a dramatic, inverse pattern of RNA translation in striatal microglia during the initial stage of opioid withdrawal, when behavioral signs of withdrawal are dramatic. While microglia have not become classically activated, there are changes in microglia morphology that appear to be cAMP mediated and could be related to microglia-neuron interaction. Further, the induction of cAMP signaling in microglia appears to be an important and early compensatory response to opioid withdrawal. Future studies should investigate the potential therapeutic value of modulating microglia-specific mediators of these signaling events and explore the physical and interaction between neurons that occurs.

## Methods

### Animals

Experiments were performed in compliance with the Guide for the Care and Use of Laboratory Animals (NIH, 1985; Publication 865–23) and were approved by the Institutional Animal Care and Use Committee, University of Washington.

### Experimental design

The experimental groups consisted of both male and female mice generated by crossing tamoxifen-inducible CX3CR1-CreERT2 hemizygous mice with homozygous floxed RiboTag mice (all on the C57Bl6 background). The experiment was performed as a 2×2 treatment design (n=6/group) with all mice receiving tamoxifen for seven days at six weeks. At ~11 weeks of age the mice received either morphine or saline for five days, on a twice daily schedule with the morphine dose escalating as follows: 20 mg/kg *ip* BID (day 1), 40 mg/kg *ip* BID (day 2), 60 mg/kg *ip* BID (day 3), 80 mg/kg *ip* BID (day 4), 100 mg/kg *ip* BID (day 5). On day 6 the mice received 100 mg/kg *ip* morphine or equivalent volume saline. Then 2 hours later they received either naloxone (1 mg/kg *sc*) or saline and the mice were video recorded for 30 minutes from above for automated scoring of withdrawal behaviors in individual cages using Ethovision (Noldus, Wageningen NL) and custom MATLAB scripts (Mathworks, Natick MA). Three variables were calculated to behavioral analysis. Distance, defined as total distance moved in cm over 30 minutes; Contraction, defined as total time spent occupying less than 65% total body length (occurs while hunching); Immobility, defined as total time spent moving less than 2cm per second. Two-way ANOVA were performed in MATLAB, followed by ‘Tukey-Kramer’ corrected contrasts. Animals were sacrificed at 2 hours after naloxone/vehicle injection. Two animals in the saline-saline (SS) group were lost during tamoxifen gavage, 1 animal from the morphine-saline (MS) was lost to an inguinal hernia, and video from 2 saline-naloxone (SN) animals were corrupted, which is reflected in figure 2. Only 4 samples from each group (2 males, 2 females) were planned for sequencing, which was still available in every group. Code used for figure generation and statistics are available in the Supplementary Code and Data.

### RiboTag RNA Isolation and Quantification

The striatal punches were homogenized in supplemented homogenization buffer [HB: 500 µL per well, 50 mM Tris-HCl, 100 mM KCl, 12 mM MgCl2, 1% NP40, 1 mM DTT, 1× Protease inhibitor cocktail (Sigma-Aldrich), 200 U/mL RNAsin (Promega, Madison, WI), 100 µg/mL cyclohexamide (Sigma-Aldrich), 1 mg/ mL heparin (APP Pharmaceuticals, Lake Zurich, IL)]. Samples were centrifuged at 4°C at 11,934 × g for 10 min, and supernatant was collected. 50µL aliquot was added to 350µL of Supplemented RLT Lysis Buffer and stored O/N at −80 to generate the input fraction of the homogenate (Qiagen Cat. 74034, Hilden, Germany). Mouse monoclonal HA-specific antibody (2.5 µL) (HA.11, MMS-101R, ascites fluid; Covance, Princeton, NJ) was added to the remaining supernatant, and RiboTag-IP fractions were rotated at 4°C for 4 h. Protein A/G magnetic beads (100 µL) (Promega) were washed with HB prior to addition to the RiboTag-IP fraction and were rotated at 4°C overnight. The next day, RiboTag-IP fractions were placed on DynaMag-2 magnet (Life Technologies), and the bead pellet was washed 3 times for 5 min with high salt buffer (HSB; 50 mM Tris, 300 mM KCl, 12 mM MgCl2, 1% NP40, 1 mM DTT, and 100 µg/mL cyclohexamide). After the final wash, HSB was removed and beads were re-suspended in 400 µL supplemented RLT buffer (10 µL β-mercaptoethanol/10 mL RLT Buffer) from the RNeasy Plus Micro Kit (Qiagen, Hilden, Germany) and vortexed vigorously. RLT buffer containing the immunoprecipitated RNA was removed from magnetic beads prior to RNA purification using RNeasy Micro Plus Kit according to the manufacturer’s protocol. RNA from both IP and input fractions were isolated using RNeasy Plus Micro Kit and eluted with 15µl of water, with typical yields of 50 ng/individual sample. RNA concentration measured using Quant-iT RiboGreen RNA Assay (ThermoFisher Cat. R11490, Waltham, MA). RNA integrity was measured using High Sensitivity RNA ScreenTape on a 2200 TapeStation (Agilent Technologies, Santa Clara, CA) by the Fred Hutchinson Cancer Research Center Genomics Core Facility, RIN 7-9 for all samples.

### RNA-seq Library Preparation and qPCR

RNA-seq libraries were prepared using SMARTer Stranded Total RNA-Seq Kit v2 – Pico Input Mammalian (Takara Bio USA, Inc. Cat. 635007, Mountain View, CA). 10ng of RNA or average equivalent volumes of “no Tamoxifen” control samples were used to generate the negative control samples. RNA-seq libraries were submitted to Northwest Genomics Center at University of Washington (Seattle, WA) where library quality control was measured using a BioAnalyzer, library concentrations were measured using Qubit dsDNA HS Assay Kit (ThermoFisher), and then samples were normalized and pooled prior to cluster generation on HiSeq High Output for paired-end reads. RNA-seq libraries were sequenced on the HiSeq4000, Paired-end 75bp to sufficient read depth with PhiX spike-in controls (7%) (Illumina San Diego, CA).

### Transcript Quantification and Quality Control

Raw fastq files were processed using multiple tools through the Galaxy platform^49^. Fastq files were inspected for quality using FastQC (Galaxy Version 0.7.0), and then passed to Salmon^50^ (Galaxy Version 0.8.2) for quantification of transcripts. The Salmon index was built using the protein coding transcriptome ftp://ftp.ensembl.org/pub/release-91/fasta/mus_musculus/cdna/Mus_musculus.GRCm38.cdna.all.fa.gz.

### Non-Specific Immunoprecipitated RNA Subtraction

An issue inherent to all immunoprecipitation (IP) procedures, including RiboTag and TRAP, non-specific precipitation. Although this RNA represents a small fraction of the IP sample, large changes in the whole tissue transcript pool (Input) could lead to small but detectable changes in the IP sample. To combat this issue and improve the reliability of RiboTag-Seq, we’ve developed novel computational method for subtraction of non-specific RNA counts from the microglia specific RiboTag sample. The only downside to this approach is that for all IP samples sequenced, the corresponding Input sample must be sequenced. First, an IP is run on equivalent tissue from animals that do not express RiboTag. RNA quantity from these samples are then quantified (mean ~6ng) along with the true IP samples (mean ~50ng). Non-specific RNA contamination is a random sampling of the Input, and the proportion of the IP sample that is contaminated is calculated from the true RNA quantity calculated for each sample.

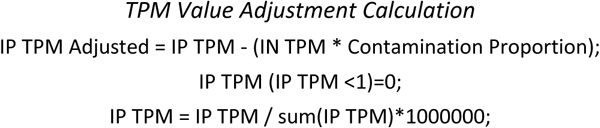

While relatively simple, this subtraction ensures that large changes in the Input sample aren’t reflected in the IP sample. For example, if an IP sample were 10% contaminated with Input RNA, a 100-fold change in the Input RNA could be captured as a 10-fold change in the IP sample. We have extensively modeled the consequences of this method of contamination subtraction and determined that it improves the reliability and interpretability of cell type specific IP, even if the level of contamination is over or underestimated (Supplementary Method 1). All IP data presented in main manuscript has undergone this adjustment, but we have also run all bioinformatics on the raw IP files, which are available in the Supplementary Code and Data. The adjusted bioinformatics are labeled “IP Adjusted” while the raw files are labeled “IP”.

### Differential Expression Analysis

Differential gene expression was calculated using DESeq2^51^ (Galaxy Version 2.11.39). All Salmon and DESeq2 settings were left default and our analysis pipeline is archived on our http://172.25.144.218:8080/u/kcgalaxy/h/morphine-rna-seq-run-2. Differential expression analysis was performed on the IP and Input samples. To determine microglia specific gene enrichment, all IP samples were compared to all Input samples. For all other test of differential expression, IP samples were compared to IP samples, and Input to Input. To determine the effects of naloxone alone, SN mice were compared to SS mice. To determine the effects of escalating morphine, MS mice were compared to SS mice. To determine the effects of withdrawal, MN mice were compared to MS mice. For all comparisons, a positive Wald statistic means that a gene is expressed more in the first group as compared to the second group. An FDR of q = 0.10 was used as the threshold for differential expression throughout the manuscript.

### Gene Set Enrichment Analysis

Wald statics generated by DeSeq2 were used as the ranking variable for gene set enrichment analysis. The Wald statistic has a benefit over log2(fold change) as it incorporates an estimate of variance, and ultimately is used to determine significance in differential expression analysis. All genes with reliable statistical comparisons (those not filtered by DeSeq2) were entered into WebGestalt 2019^52^ and GSEA was run all pertinent comparisons (enrichment, IP morphine, IP withdrawal, Input morphine, and Input Withdrawal). Gene sets analyzed include GO: Biological Process, GO: Molecular Function, Keeg, Reactome, Wiki Pathway, Transcription Factor Targets, and MicroRNA Targets. All advanced parameters were left default except for significance level, which was set to FDR = 0.05.

### Weighted Gene Co-Expression Network Analysis

Topological overlap matrix for IP and Input samples were generated, and module clustering was accomplished using the WGCNA ^27^ package for R ^53^. Briefly, the TPM matrix for each group was filtered to remove zero-variance genes, and a signed adjacency matrix was generated using “bicor” as the correlation function. From this a signed a dissimilarity topological overlap matrix was generated. Finally, module membership was assigned using a dynamic tree cut.

Post processing of the topological overlap matrix was completed with a custom MATLAB class structure that we are releasing alongside this manuscript. The @WGCNA class for MATLAB allows simple object-based analysis of complex WGCNA data. Briefly, a topological overlap matrix, module membership file, and DeSeq (or comparable) files can be loaded into a single “W” object. The WGCNA class allows for calculation and visualization of module eigen genes, merging of modules, and analyses of module membership and differential expression. Further, the WGCNA class takes full advantage of MATLAB’s powerful graph and network analysis functions. “W” objects can be used to generate complete genome minimum spanning trees with module membership or differential expression values as node markers, 2D or 3D graphs from individual modules or individual genes, with flexible settings edge pruning, node appearance, and edge appearance, etc. The WGCNA class structure can be accessed on https://Github.com/DrCoffey/WGCNA and the code used to generate all of the WGCNA figures in this manuscript is available in the Supplementary Code and Data.

### Inhibition of cAMP signaling in Microglia via hM_4_Di

Experimental animals were generated by crossing mice that are hemizygous for tamoxifen-inducible CX3CR1-Cre^ERT2^ with mice that are hemizygous for floxed hM_4_Di. Tamoxifen induction of CRE was performed in the same way as the RiboTag experiment. 2 groups of CX3CR1-hM_4_Di and 1 group of CX3CR1-Cre^ERT2^ controls with 3 males and 3 females per group were used. All animals receiving escalating morphine as described in above; 2 hrs after the final morphine injection (100 mg/kg *ip*) the mice either received CNO (3 mg/kg *ip*) or 2% DMSO vehicle 20 minutes prior to naloxone (1 mg/kg *sc*). Withdrawal was automatically scored using the same custom MATLAB script described above.

### Immunohistochemistry and Image Analysis for Iba1 Staining

The experimental groups consisted of both male and female C57BL/6 mice. The experiment was performed as a 2×2 treatment design (n=4/group). Mice underwent identical morphine administration and naloxone injections as the RiboTag-Seq experiment. Two hours following naloxone injections, mice were deeply anesthetized with an intraperitoneal injection of sodium pentobarbital and phenytoin sodium and perfused transcardially with 100 ml of 0.1 M phosphate-buffered saline (PBS), followed by 250 ml of cold 4% paraformaldehyde (pH 7.4). Brains were dissected, post fixed in 4% paraformaldehyde overnight at 4°C and transferred to 30% sucrose in PBS. Sections (40 µm) across the rostro-caudal axis of the striatum were collected on a sliding freezing microtome.

For Iba1 immunohistochemistry, free floating sections were rinsed in 1X PBS (3X, 10 minutes), permeabilized with 0.5% Triton PBS, and blocked with 4% BSA in 0.3% Triton PBS at room temperature. Sections were incubated with primary anti-Iba1 rabbit antibody diluted 1:500 (FUJIFILM Wako; #019-19741) in 4% BSA and 0.3% Triton PBS for 18h at 4°C. Sections were then rinsed with PBS (3X, 10 minutes), incubated with 1: 400 dilution of Alexa Fluor 488 Goat anti-rabbit secondary antibody (ThermoFisher; #A-11008) for sixty minutes at room temperature, washed with PBS (3X 10 minutes), mounted on charged slides, and cover slipped with Pro-Long Gold mounting medium (ThermoFisher; #P36935). Confocal stacks (30µm @40x) of the striatum (AP 0mm; ML 2mm; DV −3mm) were acquired on a Leica SP8X confocal microscope. Individual microglia reconstruction and quantification was performed using 3DMorph software ^29^. Images were processed in a blind batch, and all image analysis settings remained identical. The settings and code used to generate these analyses are available in the Supplementary Code and Data.

## Supporting information

Supplementary Modeling

## Code and Data Availability

All code and data used to generate this manuscript are available for direct download. All RNA-Seq files and the RNA-Seq pipeline are available on our Galaxy Server. All code and data used for figure generation are available in the Supplementary Code and Data. The topological overlap Matrices are also hosted on the Galaxy server because of file size limitations associated with GitHub.

## Acknowledgments

Supported by R21 DA044757 and T32 DA007278.

## Author Contributions

Kevin Coffey performed the experiment, analyzed the data, and wrote the manuscript. Atom Lesiak performed the experiment and analyzed the data. Russell Marx analyzed the data. Gwenn Garden provided animals, assisted in data interpretation, and wrote the manuscript. John Neumaier designed the experiment and wrote the manuscript.

---

**Supplementary Figure 1.**
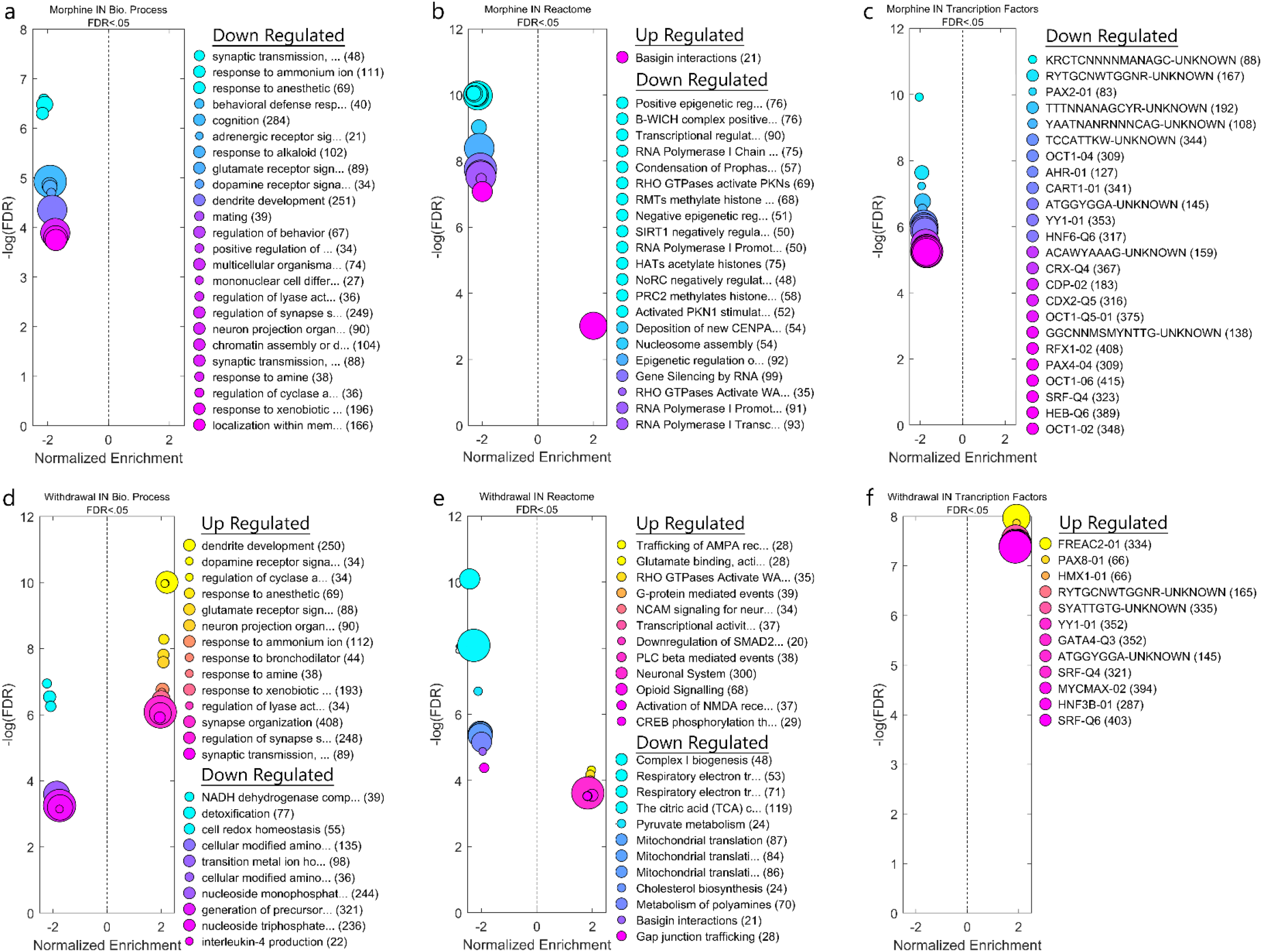
Gene Set Enrichment Analysis (Input) GSEA reveals many annotated gene sets that are significantly upregulated or downregulated in the Input samples. **a,b,c**, For example, morphine tolerance upregulates basigin interactions, and downregulates synaptic transmission genes and genes associated with the transcription factor PAX2. **d,e,f**, On the other hand, withdrawal upregulates dendritic development genes, trafficking of AMPA receptors, and genes associated to the immune related transcription factor PAX-8, and downregulation of detoxification genes, genes associated to pyruvate metabolism.

**Supplementary Figure 2.**
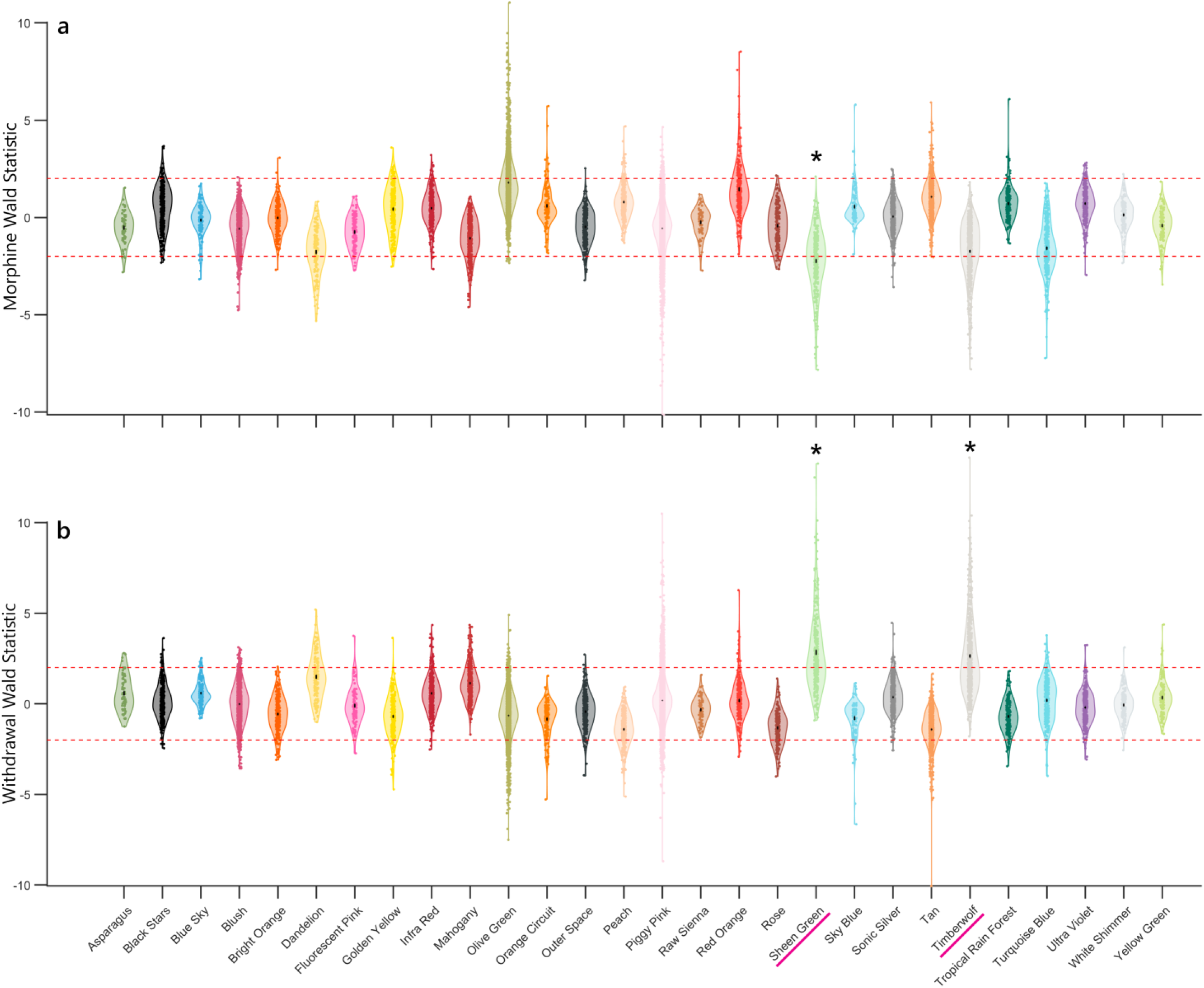
WGCNA Module Significance. Most gene modules are unrelated to morphine tolerance and withdrawal. However, two gene modules, Sheen Green and Timberwolf, are on average **a**, downregulated during morphine tolerance, and **b**, upregulated in withdrawal.

