## Supplementary Table 1 for "RiboTag-Seq Reveals a Compensatory cAMP Responsive Gene Network in Striatal Microglia Induced by Morphine Withdrawal"

| **Gene Name** | **Module** | **Centrality** | **Full Name** | **Description** |
| --- | --- | --- | --- | --- |
| Smpd3 | Sheen Green | 0.004772383 | Sphingomyelin phosphodiesterase 3 | Catalyzes the hydrolysis of sphingomyelin to form ceramide and phosphocholine. Ceramide mediates numerous cellular functions, such as apoptosis and growth arrest, and can regulate these 2 cellular events independently. |
| Cacna2d3 | Sheen Green | 0.004760259 | Calcium Channel, Voltage-Dependent, Alpha 2/Delta Subunit 3 | The alpha-2/delta subunit of voltage-dependent calcium channels regulates calcium current density and activation/inactivation kinetics of the calcium channel. |
| Phactr1 | Sheen Green | 0.004750367 | Phosphatase and Actin Regulator 1 | Binds actin monomers (G actin) and plays a role in multiple processes including the regulation of actin cytoskeleton dynamics, actin stress fibers formation, cell motility and survival, formation of tubules by endothelial cells, and regulation of PPP1CA activity. Involved in the regulation of cortical neuron migration and dendrite arborization. |
| Syndig1l | Sheen Green | 0.004750238 | Synapse Differentiation Inducing 1 Like | May regulate AMPA receptor content at nascent synapses and have a role in postsynaptic development and maturation. Expressed highly in human microglia. |
| Arhgef9 | Sheen Green | 0.004745868 | Cdc42 Guanine Nucleotide Exchange Factor 9 | The protein encoded by this gene is a Rho-like GTPase that switches between the active (GTP-bound) state and inactive (GDP-bound) state to regulate CDC42 and other genes. This brain-specific protein also acts as an adaptor protein for the recruitment of gephyrin and together these proteins facilitate receptor recruitment in GABAergic and glycinergic synapses. |
| Pdyn | Sheen Green | 0.00474121 | Prodynorphin | The protein encoded by this gene is a preproprotein that is proteolytically processed to form the secreted opioid peptides beta-neoendorphin, dynorphin, leu-enkephalin, rimorphin, and leumorphin. These peptides are ligands for the kappa-type of opioid receptor. |
| Jcad | Sheen Green | 0.004740184 | Junctional Cadherin 5 Associated | This gene encodes an endothelial cell-to-cell junction protein. Also highly expressed in mouse brain tissue. |
| Pde10a | Sheen Green | 0.004735912 | Phosphodiesterase 10A | The protein encoded by this gene belongs to the cyclic nucleotide phosphodiesterase family. It plays a role in signal transduction by regulating the intracellular concentration of cyclic nucleotides. This protein can hydrolyze both cAMP and cGMP to the corresponding nucleoside 5' monophosphate, but has higher affinity for cAMP, and is more efficient with cAMP as substrate. |
| Itpka | Sheen Green | 0.004729103 | Inositol-Trisphosphate 3-Kinase A | Regulates inositol phosphate metabolism by phosphorylation of second messenger inositol 1,4,5-trisphosphate to Ins(1,3,4,5)P4. It is also a substrate for the cyclic AMP-dependent protein kinase, calcium/calmodulin- dependent protein kinase II, and protein kinase C in vitro. |
| Gldc | Sheen Green | 0.004728291 | glycine decarboxylase | Degradation of glycine is brought about by the glycine cleavage system, which is composed of four mitochondrial protein components: P protein (a pyridoxal phosphate-dependent glycine decarboxylase), H protein (a lipoic acid-containing protein), T protein (a tetrahydrofolate-requiring enzyme), and L protein (a lipoamide dehydrogenase). |
