## Supplementary Table 2 for "RiboTag-Seq Reveals a Compensatory cAMP Responsive Gene Network in Striatal Microglia Induced by Morphine Withdrawal"

| **Name** | **module** | **Centrality** | **Full Name** | **Description** |
| --- | --- | --- | --- | --- |
| Cdk17 | Timberwolf | 0.002105623 | Cyclin Dependent Kinase 17 | The protein encoded by this gene belongs to the cdc2/cdkx subfamily of the ser/thr family of protein kinases. It has similarity to a rat protein that is thought to play a role in terminally differentiated neurons. |
| Rragd | Timberwolf | 0.002104822 | Ras Related GTP Binding D | RRAGD is a monomeric guanine nucleotide-binding protein, or G protein. By binding GTP or GDP, small G proteins act as molecular switches in numerous cell processes and signaling pathways. |
| Rasgef1a | Timberwolf | 0.002103086 | RasGEF Domain Family Member 1A | Guanine nucleotide exchange factor (GEF) with specificity for RAP2A, KRAS, HRAS, and NRAS (in vitro). Plays a role in cell migration. Among its related pathways are RET signaling and Cytokine Signaling in Immune system. |
| Kctd1 | Timberwolf | 0.002102006 | Potassium Channel Tetramerization Domain Containing 1 | The encoded protein negatively regulates the AP-2 family of transcription factors and the Wnt signaling pathway. Among its related pathways are Activation of cAMP-Dependent PKA and Sweet Taste Signaling. Gene Ontology (GO) annotations related to this gene include transcription factor binding and transcription corepressor activity. |
| Prkcb | Timberwolf | 0.002101362 | Protein Kinase C Beta | Protein kinase C (PKC) is a family of serine- and threonine-specific protein kinases that can be activated by calcium and second messenger diacylglycerol. PKC family members phosphorylate a wide variety of protein targets and are known to be involved in diverse cellular signaling pathways. |
| Ppp3r1 | Timberwolf | 0.002101225 | Protein Phosphatase 2B Regulatory Subunit 1 | Regulatory subunit of calcineurin, a calcium-dependent, calmodulin stimulated protein phosphatase. Confers calcium sensitivity. Among its related pathways are CLEC7A (Dectin-1) signaling and Wnt Signaling Pathways. |
| Grin1 | Timberwolf | 0.002100792 | Glutamate Receptor Ionotropic, NMDA 1 | The protein encoded by this gene is a critical subunit of N-methyl-D-aspartate receptors, members of the glutamate receptor channel superfamily which are heteromeric protein complexes with multiple subunits arranged to form a ligand-gated ion channel, and play a key role in the plasticity of synapses. |
| Gnal | Timberwolf | 0.002100503 | G Protein Subunit Alpha L | Guanine nucleotide-binding proteins (G proteins) are involved as modulators or transducers in various transmembrane signaling systems. G(olf) alpha mediates signal transduction within the olfactory neuroepithelium and the basal ganglia. |
| Ppp2r1a | Timberwolf | 0.002098965 | Protein Phosphatase 2 Scaffold Subunit Aalpha | The PR65 subunit of protein phosphatase 2A serves as a scaffolding molecule to coordinate the assembly of the catalytic subunit and a variable regulatory B subunit. Upon interaction with GNA12 promotes dephosphorylation of microtubule associated protein TAU/MAPT |
| Ccsap | Timberwolf | 0.002097758 | Centriole, Cilia and Spindle Associated Protein | Plays a role in microtubule (MT) stabilization and this stabilization involves the maintenance of NUMA1 at the spindle poles. Colocalizes with polyglutamylated MTs to promote MT stabilization and regulate bipolar spindle formation in mitosis. |
