## Supplementary Modeling for "RiboTag-Seq Reveals a Compensatory cAMP Responsive Gene Network in Striatal Microglia Induced by Morphine Withdrawal"

Untitled 

Supplementary Method: Removing Input Contamination from RiboTag RNA-Seq data

One major concern with Ribotag-RNA-Seq data is that non-specific RNA can contaminate immunoprecipitated RNA before sequencing. Even though the quantity of non-specific RNA is generally small, large changes in gene expression in the "whole tissue" transcriptome (Input) could manifest as small but significant changes in the RiboTag (IP) sample. However, by running proper negative controls, and sequencing both the Input and IP samples, it is possible to mathematicall deontaminate the IP sample, providing an extremely pure representation of RIboTag RNA.

Bellow we use a combination of actual data and modeling to show how this technique can be applied, and the efficay of noise removal even given imperfect assumption about the quantity of contamination. To our knowlege, this is the first method to systematically model and remove Input contimnation from RiboTag RNA-Seq data.

Understanding the Level of Contamination

Ribotag in the present study was expressed in microglia by crossing floxed RiboTag mice with tamoxifen inducible cx3cr1-CRE mice. By ommiting tamoxifen administration from a 2 animals and running the complete RiboTag protocol in paralled with the true RiboTag animals, we can get a good estimate of non-specific RNA pulldown from our procedures. All RNA is reported in nanograms. In the present study we estimate a range of contamination from 6.6-15.9% with an average of 12%.

RiboTag\_RNA=([54.65 60.76 52.94 44.24 80.60 90.52...

80.97 68.37 37.43 44.03 38.49 56.53...

33.01 38.58 44.74 46.11 42.19]); % Actual RNA Yield

Negative\_Control\_RNA=([6.56 5.39]); % Actual RNA Yield

Estimated\_Contamination=mean(Negative\_Control\_RNA)./RiboTag\_RNA;

Mean\_Contamination=mean(Estimated\_Contamination);

figure('Position',[1 1 300 300]);

g=gramm('x',zeros(17,1)','y',Estimated\_Contamination);

g.stat\_violin('normalization','width','fill','transparent');

g.geom\_jitter('width',.1);

g.stat\_summary('geom',{'black\_errorbar'},'type','sem');

g.axe\_property('XLim',[-1 1],'YLim',[0 .20],'tickdir','out');

g.set\_names('x',[],'y','Estimated Contamination (Proportion)','color','Gene Length');

g.set\_title('Fig. 1 - Input Contamination');

g.draw;

Generating "True" Gene Counts for Contamination Modeling

First we generate a model of "true" gene quantity for the Input and IP samples. These values represent the underlying "true" quantity of each gene, and are generated using a gaussian distribution with a mean of 4. Gene quantities in the IP and Input sample are not constrained and may differ greatly for any given gene, as is the case with genes in different cell types.

number\_of\_genes = 20000;

gene\_quantity\_IN = randg(4,number\_of\_genes,1);

gene\_quantity\_IP = randg(4,number\_of\_genes,1);

figure('Position',[1 1 400 300]);

histogram(gene\_quantity\_IP);

box off;

xlabel('Gene Quantity');

ylabel('Number of Genes');

title("Fig. 2 - 'True' Gene Quantity Histogram (IP)");

figure('Position',[1 1 400 300]);

scatter(gene\_quantity\_IP,gene\_quantity\_IN,4,"black","filled");

box off;

xlabel('IP Gene Quantity');

ylabel('Input Gene Quantity');

title("Fig. 3 - IP x Input Relationship");

Generating Simulated RNA-Seq Reads and TPM

RNA-Seq is based around technology that reads a finite amount of gene fragments. One consequence of this method is that the final data are a proprtional representation of the "true" gene counts, where longer genes are more likely to be counted. In order to control for these phenomenon, specialized units have been developed such as transcripts per million reads (TPM). When calculating TPM, you normalize for gene length first, and then normalize for sequencing depth second. The sum of all TPMs in each sample are the same (1e6). This makes it easier to compare the proportion of reads that mapped to a gene in each sample, and possible to use Inupt TPM to modify IP TPM.

Below we model the RNA-Sequencing machines bias for long genes, and show how calculating TPM can correct this bias and produce values that nearly perfectly match the "true" underlying gene count per million reads (CPM).

number\_of\_reads = 1e7; % Read depth of our modeled sequencing run

gene\_length = randg(3,number\_of\_genes,1); % Random gene length from gamma (mean=3)

% IN

read\_probability\_IN = gene\_quantity\_IN .\* gene\_length;

% Normalized read probability

read\_probability\_IN = read\_probability\_IN ./ sum(read\_probability\_IN);

% Generate 1e7 sequencing counts based on the read probability of each gene

read\_count\_IN = poissrnd(number\_of\_reads \* read\_probability\_IN);

TPM\_IN = calculateTPM(read\_count\_IN, gene\_length); % Calculate TMP

% IP

read\_probability\_IP = gene\_quantity\_IP .\* gene\_length;

% Normalized read probability

read\_probability\_IP = read\_probability\_IP ./ sum(read\_probability\_IP);

% Generate 1e7 sequencing counts based on the read probability of each genes

read\_count\_IP = poissrnd(number\_of\_reads \* read\_probability\_IP);

TPM\_IP = calculateTPM(read\_count\_IP, gene\_length); % Calculate TMP

figure('Position',[1 1 500 400]);

g = gramm('x',gene\_quantity\_IP,'y',read\_count\_IP, 'color', gene\_length);

g.set\_continuous\_color;

g.geom\_point;

g.set\_names('x','Gene quantity','y','Read count','color','Gene Length');

g.set\_title('Fig. 4 - Simulated Reads (IP)');

g.axe\_property('XLim',[0 20],'YLim',[0 5000],'tickdir','out');

g.draw;

figure('Position',[1 1 500 400]);

g = gramm('x',1e6 \* gene\_quantity\_IP / sum(gene\_quantity\_IP),'y',TPM\_IP, 'color', gene\_length);

g.set\_continuous\_color;

g.geom\_point;

g.set\_names('x','Real CPM','y','Observed TPM','color','Gene Length');

g.set\_title('Fig. 5 - TPM (IP)');

g.axe\_property('XLim',[0 250],'YLim',[0 250],'tickdir','out');

g.draw;

r\_squared = corr(TPM\_IP,gene\_quantity\_IP)^2

r\_squared = 0.9881

Contaminating the IP Sample with Gene Counts from the Input

Now we can contaminate our "true" IP gene counts with gene counts from the Input sample. Below we will assume that the mean level of Input contamination from our actual samples (~12%) is mixed in with the IP genes, and see how it affects the sequencing run and our estimate of TPM.

contamination = Mean\_Contamination; % Mean observed contamination

% Contaminate the IP sample with Input genes

read\_probability\_IP\_contaminated = read\_probability\_IP\*(1-contamination) + read\_probability\_IN\*contamination;

% Generate 1e7 sequencing counts based on the read probability of each gene

read\_count\_IP\_contaminated = poissrnd(number\_of\_reads \* read\_probability\_IP\_contaminated);

TPM\_IP\_contaminated = calculateTPM(read\_count\_IP\_contaminated, gene\_length);

figure('Position',[1 1 500 400]);

g = gramm('x', 1e6 \* gene\_quantity\_IP / sum(gene\_quantity\_IP),'y', TPM\_IP\_contaminated, 'color', gene\_length);

g.geom\_point;

g.set\_names('x','Real CPM','y','Observed TPM with contamination','color','Gene Length');

g.set\_title('Fig. 6 - Contaminated TPM (IP)');

g.axe\_property('XLim',[0 250],'YLim',[0 250],'tickdir','out');

g.draw;

r\_squared = corr(TPM\_IP\_contaminated,gene\_quantity\_IP)^2

r\_squared = 0.9669

Removing Input Contamination from IP TPM

The contamination in our IP sample means that the TPMs we calculate from our modeled sequencing no longer perfectly reflect the underlying "true" gene counts. To fix this we can sequence the Input sample and simply subtract the proprtion of reads that are present due to non-specific RNA contamination.

% Subtract TPM present due to Input contamination.

TPM\_IP\_contaminated\_corrected = TPM\_IP\_contaminated - contamination\*TPM\_IN;

figure('Position',[1 1 500 400]);

g = gramm('x', 1e6 \* gene\_quantity\_IP / sum(gene\_quantity\_IP),...

'y', TPM\_IP\_contaminated\_corrected, 'color', gene\_length);

g.geom\_point;

g.set\_names('x','Real TPM','y','Observed TPM with contamination correction','color','Gene Length');

g.set\_title('Fig. 7 - Contamination Corrected TPM (IP)');

g.axe\_property('XLim',[0 250],'YLim',[0 250],'tickdir','out');

g.draw;

r\_squared = corr(TPM\_IP\_contaminated\_corrected,gene\_quantity\_IP)^2

r\_squared = 0.9847

Scalling Corrected IP TPM

Pure subtraction of TPM will lower the number of reads bellow 1e6 and some small fraction of TPM will become negative. By setting negative reads to zero, and rescalling TPM to 1e6 reads, we can restore our corrected TPM to a nearly perfect estimate of the true underlying CPM.

TPM\_IP\_contaminated\_corrected = TPM\_IP\_contaminated - contamination\*TPM\_IN ;

TPM\_IP\_contaminated\_corrected(TPM\_IP\_contaminated\_corrected<0)=0;

TPM\_IP\_contaminated\_corrected = ...

TPM\_IP\_contaminated\_corrected/sum(TPM\_IP\_contaminated\_corrected)\*1000000;

figure('Position',[1 1 500 400]);

g = gramm('x', 1e6 \* gene\_quantity\_IP / sum(gene\_quantity\_IP),...

'y', TPM\_IP\_contaminated\_corrected, 'color', gene\_length);

g.geom\_point;

g.set\_names('x','Real CPM','y','Observed TPM with contamination correction','color','Gene Length');

g.set\_title('Fig. 8 - Contamination Corrected & Scaled TPM');

g.axe\_property('XLim',[0 250],'YLim',[0 250],'tickdir','out');

g.draw;

r\_squared = corr(TPM\_IP\_contaminated\_corrected,gene\_quantity\_IP)^2

r\_squared = 0.9847

Quantifying Benefit of Correcting Given a Contamination Estimate Range

Contamination for each sample is estimated from our negative controls. This means there is a risk of over or under estimating contamination in any given sample. However, correcting for contamination anywhere within our actual range is better then no correction at all. In all RiboTag RNA-Seq experiments, large changes in the Input sample could be responsible for changes in the IP samples. Using our current experiment as an example, if morphine withdrawal causes a 100 fold increase in a particular gene in the Input, a 10% contaminated IP sample would show a 10 fold increase. Using this technique, we can reliably remove known changes in the Input samples from the IP samples, markedly improving the reliability and interpretability of our RiboTag RNA-seq results.

r\_squared = [];

contamination\_values = linspace(0, .25, 50);

for i = 1:length(contamination\_values)

TPM\_IP\_contaminated\_corrected = TPM\_IP\_contaminated - contamination\_values(i)\*TPM\_IN;

r\_squared(i) = corr(TPM\_IP\_contaminated\_corrected,gene\_quantity\_IP)^2;

end

figure('Position',[1 1 500 400]);

clear g;

g = gramm('x', contamination\_values, 'y', r\_squared);

g.geom\_line;

g.set\_names('x','Estimated contamination','y','R Squared');

g.set\_title('Fig. 9 - R^2 Inaccurate Contamination Estimates');

g.geom\_vline('xintercept',min(Estimated\_Contamination),'style','b--');

g.geom\_vline('xintercept',max(Estimated\_Contamination),'style','b--');

g.draw;

g.update('x',[0, contamination-.05],'y',[r\_squared(1)-.0003,.99],...

'label',{'No correction','Actual Contamination Range'});

New X or Y of different size given, all data from first gramm cleared

g.axe\_property('YLim',[.965 .99],'tickdir','out');

g.geom\_label;

g.draw;

TPM Calculation Function

function TPM = calculateTPM(reads, gene\_length)

TPM = reads ./ gene\_length;

TPM = 1e6 \* TPM / sum(TPM);

end

  
